# Brain-derived and in vitro-seeded alpha-synuclein fibrils exhibit distinct biophysical profiles

**DOI:** 10.1101/2023.10.04.560803

**Authors:** Selene Seoyun Lee, Livia Civitelli, Laura Parkkinen

**Affiliations:** Nuffield Department of Clinical Neurosciences, Oxford Parkinsons’ Disease Center, University of Oxford, UK

## Abstract

The alpha-synuclein (*α*Syn) seeding amplification assay (SAA) that allows the generation of disease-specific in vitro seeded fibrils (SAA fibrils) is used as a research tool to study the connection between the structure of *α*Syn fibrils, cellular seeding/spreading, and the clinico-pathological manifestations of different synucleinopathies. However, structural differences between human brain-derived and SAA *α*Syn fibrils have been recently highlighted. Here, we characterize biophysical properties of the human brain-derived *α*Syn fibrils from the brains of patients with Parkinson’s disease with and without dementia (PD, PDD), dementia with Lewy bodies (DLB), multiple system atrophy (MSA) and compare them to the ‘model’ SAA fibrils. We report that the brain-derived *α*Syn fibrils show distinct biochemical profiles, which were not replicated in the corresponding SAA fibrils. Furthermore, the brain-derived *α*Syn fibrils from all synucleinopathies displayed a mixture of- ‘straight’ and ‘twisted’ microscopic structures. However, the PD, PDD, and DLB SAA fibrils had a ‘straight’ structure, whereas MSA SAA fibrils showed a ‘twisted’ structure. Finally, the brain-derived *α*Syn fibrils from all four synucleinopathies were phosphorylated (S129). Interestingly, phosphorylated *α*Syn were carried over to the PDD and DLB SAA fibrils. Our findings demonstrate the limitation of the SAA fibrils modelling the brain-derived *α*Syn fibrils and pay attention to the necessity of deepening the understanding of the SAA fibrillation methodology.

## Introduction

Synucleinopathies are all characterized by abnormal accumulation of the protein alpha-synuclein (*α*Syn) in the brain. However, they are clinically and neuropathologically highly heterogeneous diseases with prominent disease-specific differences in the presentation of symptoms, rate of disease progression, and the brain regions and cell types vulnerable to *α*Syn deposition and neuronal death. In Parkinson’s disease (PD), Parkinson’s disease with dementia (PDD), and dementia with Lewy bodies (DLB), *α*Syn aggregation is found in the neuronal soma as Lewy bodies (LBs) and in the axons and dendrites as Lewy neurites (LNs) (Figure 1) (***Forno, 1988***; ***Spillantini et al., 1997***; ***Braak et al., 2003***). Furthermore, the astroglial accumulation of *α*Syn is a prominent but so far underdiagnosed pathology in these Lewy body disorders (***Altay et al., 2022***). The *α*Syn pathology in multiple system atrophy (MSA) is primarily found in oligodendrocytes as glial cytoplasmic inclusions (GCIs) (***Papp et al., 1989***).

**Figure 1.**
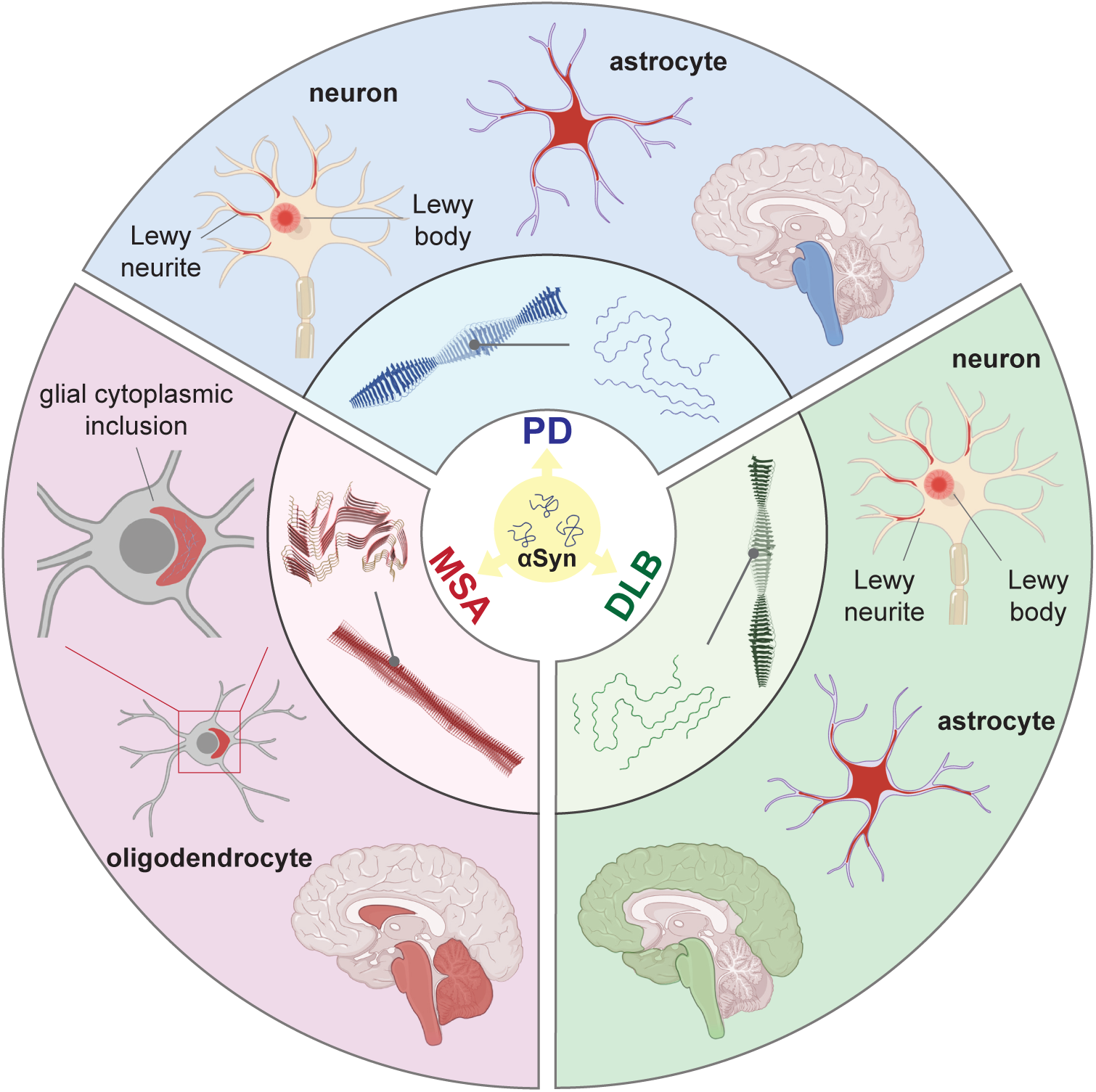
Distinct *α*Syn strains are associated with different neuropathological and clinical hallmarks of PD, DLB, and MSA. *α*Syn misfolds and aggregates into fibrils with characteristic conformations. At the atomic level, PD and DLB strains share a “Lewy fold” structure at the fibrillar core and comprise a single protofilament (***Yang et al., 2022***). MSA strains are twisted with two protofilaments intertwined, forming a different core structure to the “Lewy fold” (***Schweighauser et al., 2020***). At a cellular level, PD is characterized by significant neuronal loss at the brainstem, especially substantia nigra (SN), highlighted in blue. Lewy body (LB) and Lewy neurite (LN) accumulate in neurons. *α*Syn also accumulates in astrocytes, forming astroglial pathology. In DLB, the brainstem and neocortex are the most affected regions, highlighted in green. Here, LB and LN accumulate in neurons and astroglial pathology is also observed. In MSA, the cerebellum, basal ganglia and brainstem are the most affected, highlighted in red. Here, *α*Syn accumulates as glial cytoplasmic inclusions (GCIs) in the oligodendrocytes. The 3D structures of the *α*Syn fibrillar cores were generated using PyMOL, and the PDB structures from ***Schweighauser et al. (2020)***; ***Yang et al. (2022)***.

The mechanistic link between *α*Syn and clinico-pathological diversity of the synucleinopathies is hypothesized to be the different fibrillar ‘strains’ of *α*Syn in analogy to prion disease. A strain is generated when *α*Syn monomers fold into specific fibrillar forms with distinct conformational and biological characteristics (***Melki, 2017***). Under the disease condition, misfolded and aggregated *α*Syn recruits normal, soluble endogenous *α*Syn to aggregate, and this self-perpetuating process spreads throughout the brain-periphery axis. Several studies have generated *α*Syn fibrils in vitro using various experimental conditions to characterize the presence of different strains (***Bousset et al., 2013***; ***Peelaerts et al., 2015***; ***Li et al., 2018***; ***Guerrero-Ferreira et al., 2019***; ***Lau et al., 2020***), but their relevance to human condition remains questionable. However, some studies have also examined *α*Syn fibrils extracted from the human brain tissue of different synucleinopathies and these have demonstrated that the human-derived *α*Syn strains exhibit distinct structures (***Schweighauser et al., 2020***), infectivity and bioactivity in cells (***Woerman et al., 2018a***, ***2019***; ***Ayers et al., 2022***) and animals (***Prusiner et al., 2015***; ***Lau et al., 2020***; ***Holec et al., 2022***; ***Peng et al., 2018***). The intrinsic structure of *α*Syn proteoforms may also depend on the local conditions of the brain region where they are formed and thus may vary within a single disease entity (***Peng et al., 2018***; ***Lau et al., 2020***; ***Schweighauser et al., 2020***). Future studies must comprehensively map the distinct fingerprints of human brain-derived *α*Syn strains to the disease.

Seeding amplification assays (SAAs) have become more applicable due to their feasibility in amplifying in vitro-seeded *α*Syn fibrils (SAA fibrils) from human biosamples. SAA refers to two distinct seed amplification techniques: real-time quaking-induced conversion (RT-QuIC) and protein misfolding cyclic amplification (PMCA). Both methods incorporate a key characteristic of pathological *α*Syn, which is its ability to induce aggregation of monomeric *α*Syn into different complex conformers. When pathological *α*Syn is present in the biosample, it seeds aggregation of the recombinant *α*Syn in the reaction through repeated elongation-fragmentation cycles (***Concha-Marambio et al., 2023***). A major difference between the two methods lies in the method of aggregate fragmentation (***Coysh and Mead, 2022***). RT-QuIC uses physical shaking, and PMCA uses sonication. Also, the RT-QuIC reaction is monitored automatically in real-time using a fluorophore such as Thioflavin-T (ThT), whereas PMCA requires manual measurement. Furthermore, it should be noted that the RT-QuIC products are not infectious, while the PMCA produces infectious aggregates with specific strain fidelity (***Raymond et al., 2020***). In this study, RT-QuIC has been selectively used and referred to as ‘SAA’.

SAA, together with different biosamples such as cerebrospinal fluid (CSF) (***Fairfoul et al., 2016***; ***Poggiolini et al., 2021***), brain homogenates (***Groveman et al., 2018***; ***Sano et al., 2018***), and skin punctures (***Mammana et al., 2020***; ***Kuzkina et al., 2021***), has presented its potential as a diagnostic method but also as a research tool to amplify disease-relevant fibrils. The newly amplified SAA fibrils are assumed to encode the intrinsic properties of the original seeding fibrils, thus being representative of the disease-specific strains. Therefore, using the ‘model’ SAA fibrils, studies have shown disease-specific structural, biochemical and phenotypic differences, suggesting the presence of distinct conformational *α*Syn strains in different synucleinopathies (***Strohaker et al., 2019***; ***Shahnawaz et al., 2020***; ***Van der Perren et al., 2020***; ***Frieg et al., 2022***).

Despite such extensive strain characterisation performed with the SAA fibrils, whether the SAA fibrils are representative models of the original seed fibrils is unclear. Studies highlighted the significant structural differences between the human brain-derived and SAA fibrils (***Lövestam et al., 2021***) and seeded pathology in vivo (***Van der Perren et al., 2020***). Nevertheless, there is currently a limited understanding of whether the resulting SAA fibrils preserve other properties, including biochemical, biophysical, cellular toxicity and pathology.

Here, we aim to investigate the biophysical differences between 1) the brain-derived *α*Syn fibrils in different synucleinopathies and 2) the brain-derived *α*Syn versus the corresponding SAA fibrils. We examine the brain-derived *α*Syn fibrils extracted from the brains of patients with PD, PDD, DLB, and MSA and the brain-derived versus SAA *α*Syn fibrils and show striking differences in their biochemical profile, structure, and phosphorylation pattern. Our findings reveal further evidence supporting the molecular diversity among *α*Syn fibrils from different synucleinopathies. Also, our study highlights the limitations of the SAA fibrils to fully mirror the brain-derived *α*Syn fibrils and the need to study further the mechanism of seeding amplification and the application of SAA fibrils.

## Results

### Detection of differential *α*Syn seeding activity from brain-derived fibrils of PD, PDD, DLB, and MSA

We extracted *α*Syn fibrils from the human brain of patients with PD (*n* = 3), PDD (*n* = 3), DLB (*n* = 3), MSA (*n* = 3) and healthy controls (*n* = 3) (Table S1). The entorhinal cortex was used for PD, PDD, and DLB, and the striatum for MSA, as these are one of the main affected regions with *α*Syn pathology in each disorder. The brain-derived fibrils were diluted 1: 1,000 and used as a seed in an SAA reaction to generate in vitro amplified fibrils (SAA fibrils). PD, PDD, DLB, and MSA brain-derived *α*Syn fibrils reached the maximum ThT fluorescence within 100 h (Figure 2 A). The healthy controls did not display seeding potential except for case 2 (HC 2), which showed an increasing signal towards the end of the reaction (approximately 91 h) (Figure 2 A). The unseeded recombinant *α*Syn remained negative in all reactions (Figure 2 A). The raw data of all the replicates are presented in Figure S1.

**Figure 2.**
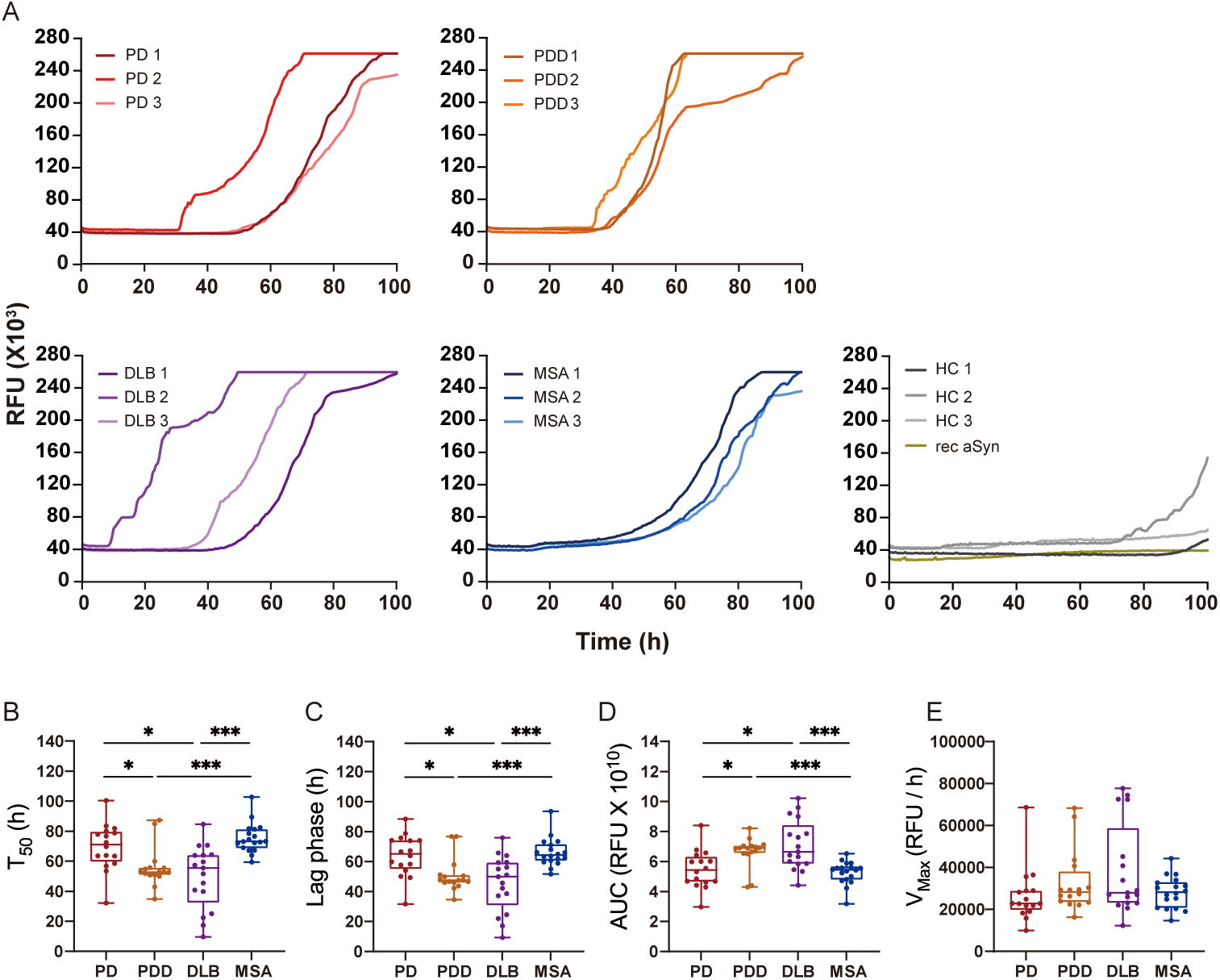
*α*Syn SAA seeded with *α*Syn fibrils from PD, PDD, DLB, and MSA brains. (**A**) SAA was performed with sarkosyl insoluble fractions of PD (*n* = 3), PDD (*n* = 3), DLB (*n* = 3), MSA (*n* = 3), and HC (*n* = 3) brains. The curves represent an average of six replicates. rec *α*Syn indicates an unseeded control reaction. (**B**) Time to reach 50% of the maximum fluorescence (*T*_50_). (**C**) The lag phase was taken at the time point where each positive reaction exceeded the threshold (RFU ≥ 5 SD). (**D**) Area under the curve (AUC). (**E**) The largest increase of fluorescence per unit of time (V_MAX_). (**B-E**) Plotted values represent the six replicates of the three cases for each disease (*N* = 18). RFU, relative fluorescence unit; SD, standard deviation. \**P* ≤ 0.05, \*\**P* ≤ 0.01, \*\*\**P* ≤ 0.005

Kinetic parameters revealed a statistical difference in the seeding activities between PD/MSA and PDD/DLB. The time to reach 50 % of the maximum fluorescence (*T*_50_) and the lag phase (time to reach five standard deviations of the minimum fluorescence) were significantly lower in PDD/DLB than in PD/MSA (Figure 2B,C). As a result, the area under the curve (AUC) was statistically higher in PDD/DLB than in PD/MSA (Figure 2D). However, the maximum rate of increase in fluorescence (*V*_MAX_) was not statistically different among the diseases (Figure 2E). Overall, the SAA kinetic parameters suggest PDD and DLB brain-derived *α*Syn fibrils have a more aggressive seeding capacity than PD and MSA.

### Brain-derived and SAA *α*Syn fibrils display distinct biochemical profiles

At the end of the SAA reaction, the SAA *α*Syn fibrils were collected by ultracentrifugation. Having established that prion strains can be characterized by distinct biochemical profiles (***Lau et al., 2020***; ***Shahnawaz et al., 2020***; ***Sano et al., 2014***), we examined this characteristic in the brain-derived and SAA *α*Syn fibrils. First, the fibrils were denatured with increasing concentrations of guanidine hydrochloride (GdnHCl, 0-5 M) and then digested with proteinase-K (PK, 1 µg/ml).

The weakest GdnHCl (1 M) treatment completely denatured the PD and MSA brain-derived *α*Syn fibrils, as they were no longer visible on the immunoblot (Figure 3). This result was consistent among all the three cases analyzed (Figure S2). Higher GdnHCl concentrations of 2 M and 3 M denatured PDD and DLB brain-derived *α*Syn fibrils, respectively (Figure 3). The results illustrate that PD and MSA brain-derived *α*Syn fibrils have weaker biochemical stability than PDD and DLB fibrils.

**Figure 3.**
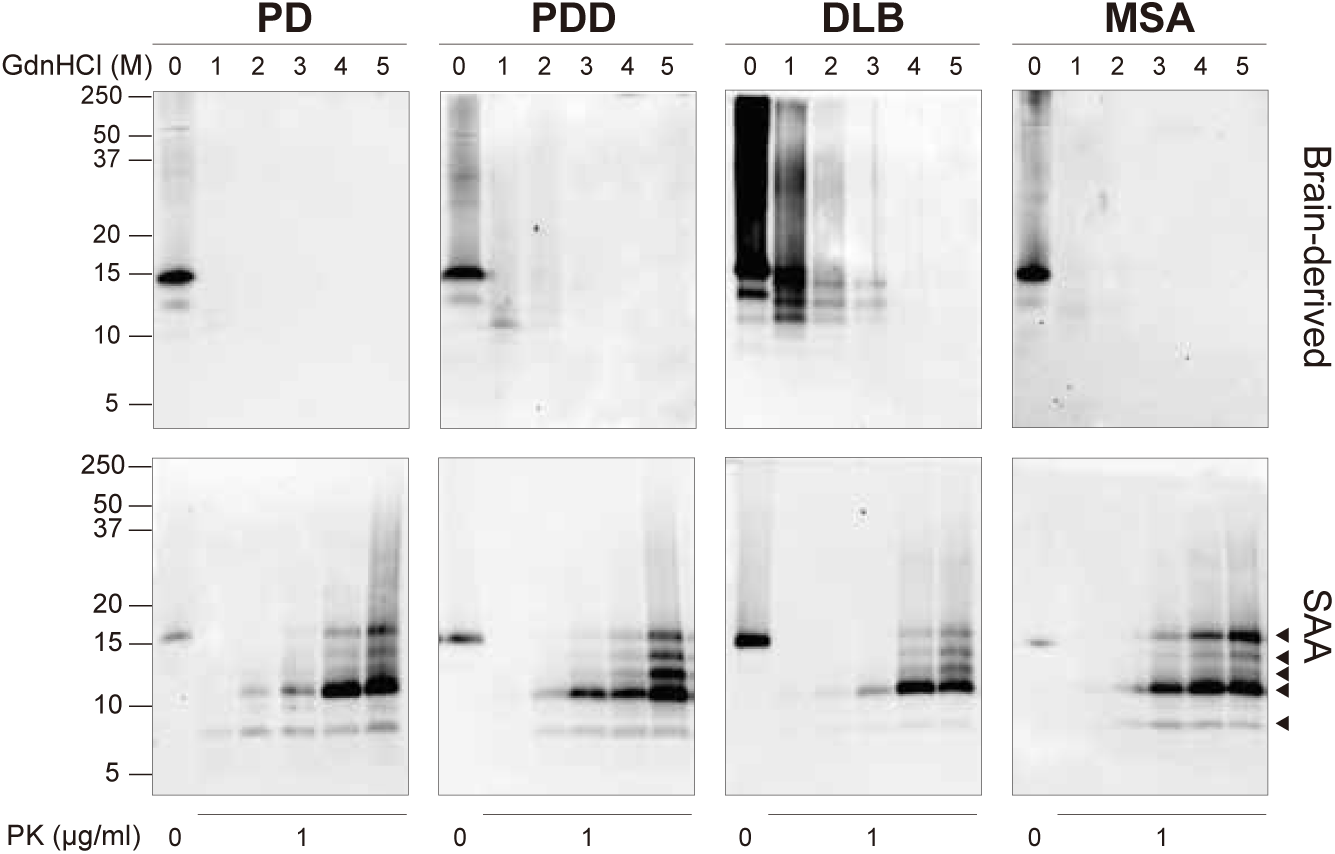
Brain-derived and SAA *α*Syn fibrils exhibited distinct biochemical profiles. Brain-derived and SAA *α*Syn fibrils were subjected to denaturation with increasing concentration of GdnHCl (0-5 M) and to PK digestion (1 µg/ml). The antibody clone 42 (BD Biosciences) was used to reveal the PK-resistant peptides. Immunoblots of one representative case from PD, PDD, DLB, and MSA are presented. The brain-derived and SAA fibrils are on the top and bottom rows, respectively. The molecular weights of the protein standards are shown in kilodaltons (kDa). Black arrows mark the five PK-resistant peptides revealed in the SAA fibrils. SAA, seeding amplification assay; GndHCl, guanidine-hydrochloride; PK, proteinase-K.

The disappearance of the bands on the immunoblot after GndHCl treatment limited the comparative study of the *α*Syn fibrils between synucleinopathies. Therefore, we repeated the experiment using PK alone to improve the visualization of the biochemical profiles. The brain-derived *α*Syn fibrils were treated with five different PK concentrations starting from 0 mg/ml, incrementing to 1 mg/ml. PD cases contained mostly monomeric *α*Syn (15 kDa) that immediately degraded at the lowest PK concentration (1 ug/ml) (Figure 4A). In contrast, in PDD, DLB, and MSA cases, high molecular weight (MW) bands above 15 kDa were observed, indicative of *α*Syn aggregates (Figure 4B-D).

**Figure 4.**
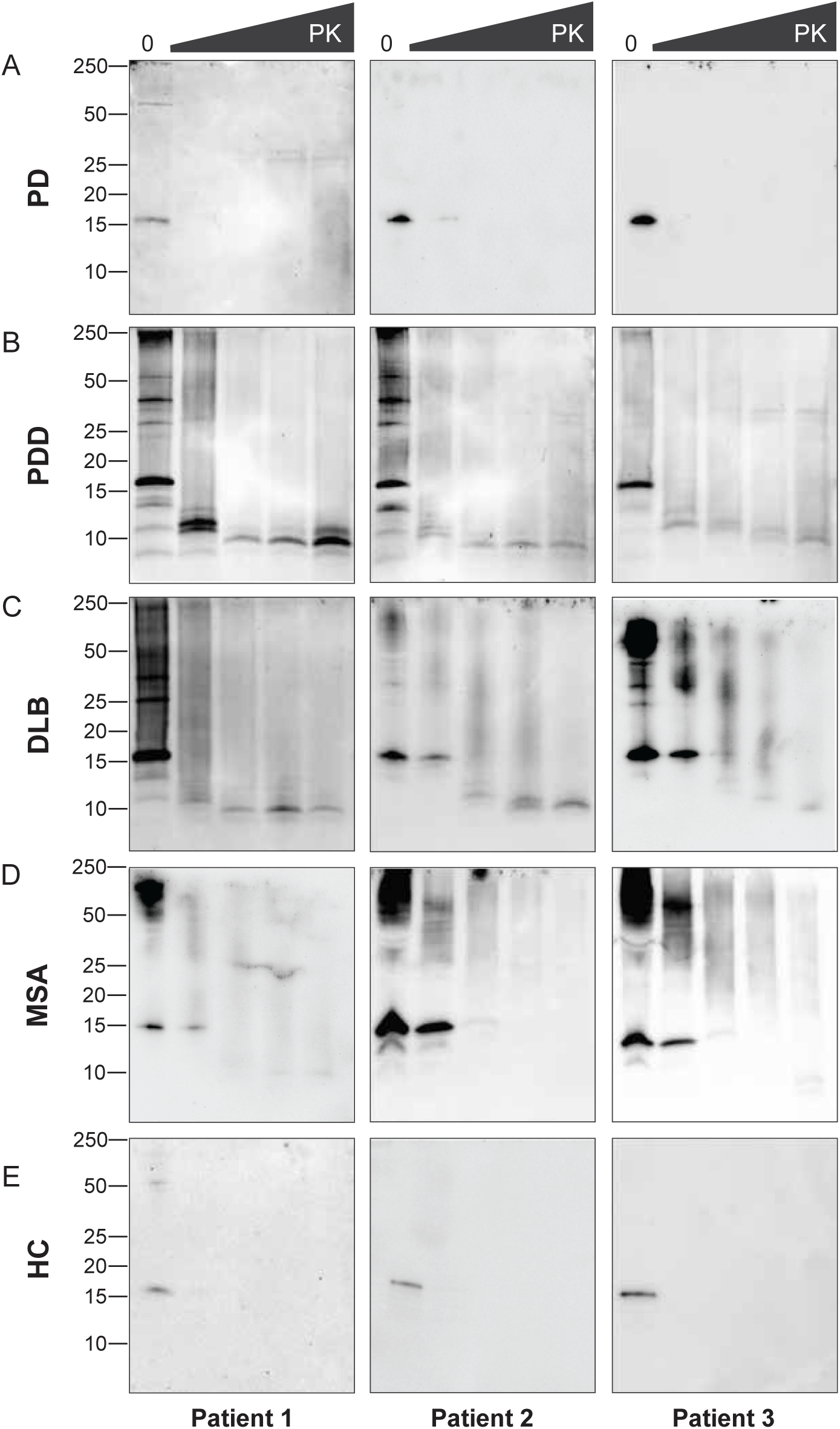
Proteinase-K degradation patterns of the brain-derived *α*Syn fibrils. The brain-derived *α*Syn fibrils from (**A**) PD, (**B**) PDD, (**C**) DLB, (**D**) MSA, and (**E**) HC were subjected to increasing concentrations of PK at 0, 0.001, 0.01, 0.1, and 1 mg/ml, represented by the escalating triangular bar. Western blot was performed using the antibody clone 42 (BD Biosciences). The molecular weights of the protein standards are shown in kilodaltons (kDa).

Furthermore, lower MW bands (13, 10, 7 kDa) were also observed. As the PK concentration increased, the PDD and DLB fibrils gradually degraded, as the high MW bands started to disappear and low MW bands below 15 kDa began to appear (Figure 4B,C). Similarly, the MSA fibrils gradually degraded as the PK concentration increased (Figure 4D). However, low MW bands below 15 kDa were less prominent than those of PDD and DLB. These low MW bands were absent in HC (Figure 4E), and PD and HC showed no difference in the PK profiles.

Although we experimented with an identical amount of total protein (10 ug), the amount of *α*Syn fibril could differ in each brain-derived sample. Using immunohistochemistry, we observed a lower load of *α*Syn pathology in PD and MSA brains than in PDD and DLB (Figure S3). In addition, the brain-derived fibrils were immunogold-labelled with anti-p*α*Syn and negatively stained for transmission electron microscopy (TEM). The samples were imaged with TEM at low magnification to observe *α*Syn fibril density and distribution. PD and MSA cases contained 1-3 *α*Syn fibrils per region of interest (ROI), whereas PDD and DLB cases contained 3-5 *α*Syn fibrils per ROI (Figure S4). Adjacent slot blots also confirmed a higher amount of *α*Syn fibrils in PDD and DLB than in PD and MSA brain-derived samples. Therefore, lower fibril concentration in the PD and MSA brains could have limited the observation of the PK-resistant peptides using immunoblotting.

The SAA fibrils revealed markedly different biochemical profiles to the brain-derived *α*Syn fibrils. We observed identical profiles among all SAA fibrils from PD, PDD, DLB, and MSA. PK-resistant bands of 7 and 10 kDa started to appear after treatment with 2 M GdnHCl (Figure 3). However, with increasing concentrations of GdnHCl, specific PK-resistant peptides appeared, consisting of five bands (7, 11, 12, 13, 14 kDa) (Figure 3). These findings suggest that harsher denaturing conditions are required to destabilize the SAA fibrils and expose PK digestion sites. We concluded that SAA *α*Syn fibrils are more stable and characterized by distinct biochemical properties than the brain-derived *α*Syn fibrils.

### Brain-derived and SAA *α*yn fibrils show structural differences

Next, we used TEM to investigate the structure of the brain-derived and SAA *α*Syn fibrils. The brain-derived *α*Syn fibrils from PD, PDD, and DLB exhibited two structures in TEM: straight and twisted (Figure 5A, Figure S5). The fibrils from MSA were predominantly straight, with the rare presence of twisted type. Therefore, we only considered straight fibrils when comparing the dimensions of the fibrils among synucleinopathies. The length of the fibrils from different synucleinopathies ranged between 68 to 885 nm. Only the PDD fibrils were significantly longer than DLB and MSA (Figure 5B). The width ranged from 7 to 21 nm, and there were significant differences between the diseases. PD had the widest fibrils, followed by MSA/DLB and PDD (Figure 5C).

**Figure 5.**
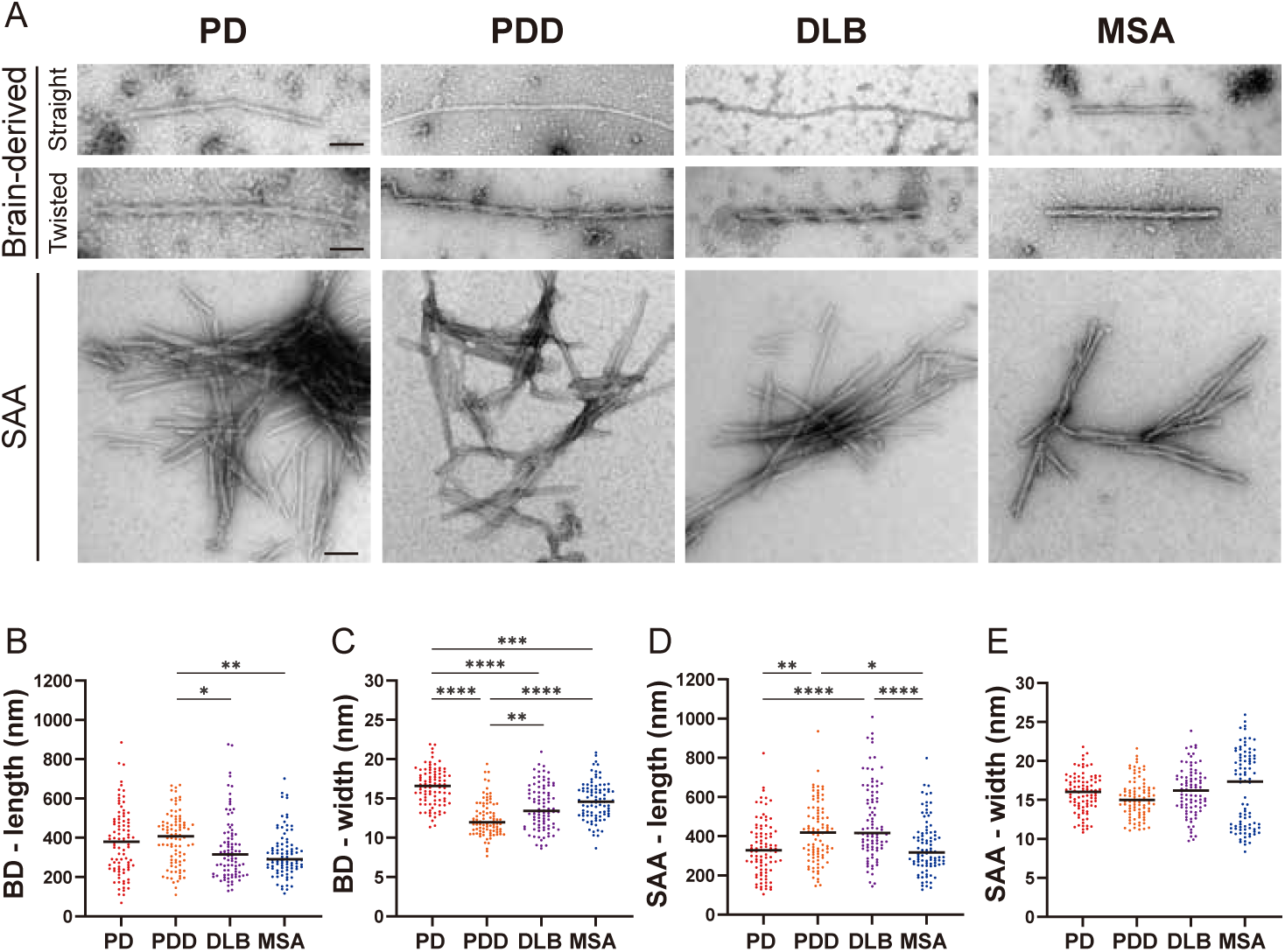
Transmission electron microscopy revealed different structures of brain-derived and SAA *α*Syn fibrils. (**A**) Electron microscope image of negatively stained brain-derived (**BD**) and SAA fibrils from PD, PDD, DLB, and MSA brains. (**B,C**) The lengths and widths of brain-derived fibrils. (**D,E**) The lengths and width of SAA fibrils. MSA SAA fibrils were twisted with alternating widths, resulting in two clusters of measurements. A total of 30 fibrils from each case were measured and plotted (*N* = 90). Scale bar = 50 nm. \**P* ≤ 0.05, \*\**P* ≤ 0.01, **** *P* ≤ 0.001.

On the other hand, the SAA fibrils demonstrated a single dominant structure. PD, PDD, and DLB SAA fibrils were all straight, and MSA SAA fibrils were all twisted (Figure 5A). Unlike the brain-derived fibrils, the SAA fibrils were highly clustered. The length of the SAA fibrils ranged from 104 to 1008 nm, and PDD/DLB SAA fibrils were significantly longer than those of PD/MSA (Figure 5D).

The width ranged from 8 to 25 nm, and the widths of the MSA SAA fibrils were distributed into two groups due to the alternating widths arising from the twisted structure. Unlike the brain-derived fibrils, the average widths of the SAA fibrils were similar between different synucleinopathies (Figure 5E).

Overall, in DLB, the SAA fibrils were significantly longer and wider than the brain-derived fibrils (Figure S6). SAA fibrils from PDD were wider than the brain-derived fibrils but not different in length. The dimensions of the brain-derived and SAA fibrils from PD and MSA were not significantly different. However, it was difficult to compare the dimensions of the MSA fibrils as the brain-derived and SAA fibrils had different morphologies. Our observations demonstrate that brain-derived and SAA *α*Syn fibrils exhibit distinct TEM structures.

### Brain-derived and SAA *α*Syn fibrils have distinct phosphorylation patterns

To confirm the identity of the fibrils imaged by TEM, we used immunogold with *α*Syn fibril conformation specific (MJFR-14) and pS129-specific (anti-p*α*Syn) antibodies. Both brain-derived and SAA fibrils from all four synucleinopathies were probed with MJFR-14, confirming that the fibrils were *α*Syn fibrils (Figure 6A). The brain-derived fibrils were also strongly labeled with anti-p*α*Syn. Both antibodies, however, did not label the twisted brain-derived fibrils from PD, PDD, and DLB (Figure 6B). The SAA fibrils from PD and MSA were not labelled with anti-p*α*Syn, but PD and MSA SAA fibrils were weakly labelled (Figure 6A).

**Figure 6.**
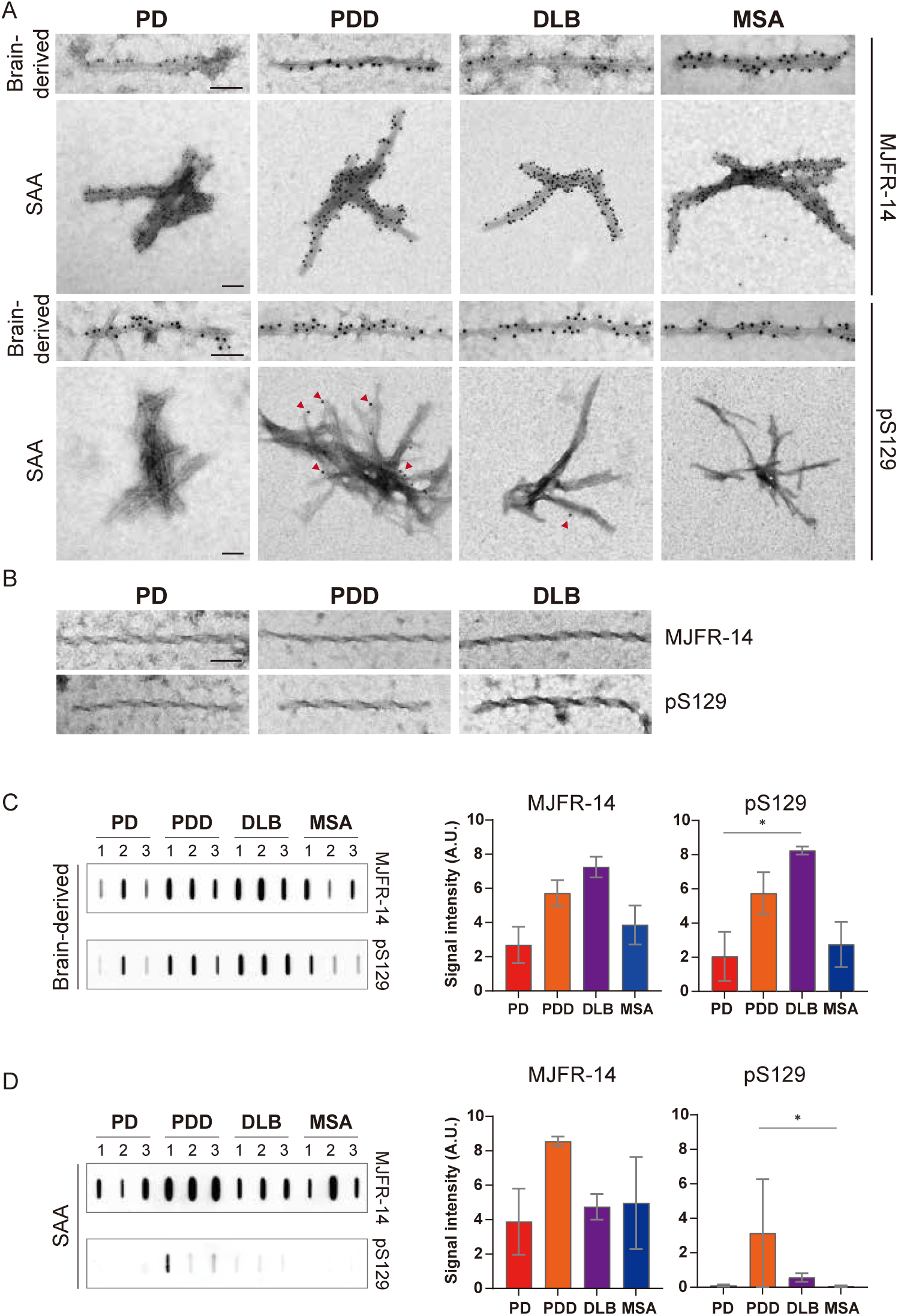
Brain-derived and SAA *α*Syn fibrils showed distinct phosphorylation patterns. (**A**) Electron microscope image of brain-derived and SAA *α*Syn fibrils from PD, PDD, DLB, and MSA brains labelled with fibril conformation-specific (MJFR-14) and anti-p*α*Syn (pS129) antibodies. (**B**) The twisted *α*Syn fibrils from PD, PDD, and DLB brains were not labelled for MJFR-14 and pS129. Scale bar = 50 nm. (**C,D**) Semi-quantification of *α*Syn fibrils and p*α*Syn confirm the different patterns of *α*Syn phosphorylation between brain-derived and SAA *α*Syn fibrils. The amount of *α*Syn fibrils and p*α*Syn in (**C**) brain-derived and (**D**) SAA fibrils was determined using a slot blot. Two µg of proteins were filtered on a nitrocellulose membrane and probed with MJFR-14 and pS129 antibodies. The semi-quantitative measurement was done by averaging the measurements of three cases from each disease. Error bars indicate ± one standard deviation (SD). * *P* ≤ 0.05.

We performed a slot blot using identical antibodies to confirm the immunogold-labelling results. On the slot blot, DLB brain-derived *α*Syn fibrils showed the strongest signal with MJFR-14 and anti-p*α*Syn, followed by PDD, MSA, and PD (Figure 6C). Interestingly, the anti-p*α*Syn signal from the SAA fibrils was markedly different from the brain-derived fibrils. In parallel to the anti-p*α*Syn immuno-gold results (Figure 6A), PDD SAA fibrils showed the highest anti-p*α*Syn signal (Figure 6D). The DLB SAA fibrils were also weakly phosphorylated, while those of PD and MSA were negative. Considering the absence of phosphorylated recombinant monomers in our SAA reaction, the phosphorylated *α*Syn must have transmitted from the seed brain-derived fibrils. Our results demonstrate that the brain-derived and SAA *α*Syn fibrils display different patterns of S129 phosphorylation.

## Discussion

### Strain-like properties of the brain-derived *α*Syn fibrils from different synucleinopathies

Findings in this study provide evidence of distinct ‘strain-like’ properties of the brain-derived *α*Syn fibrils from different synucleinopathies. First, faster seeding kinetics were observed in the brain-derived *α*Syn fibrils from PDD and DLB than those from PD and MSA, suggesting a distinct ‘strain’ characterized by an aggressive seeding capacity. In contrast, those from PD and MSA are discrete ‘strains’ with milder seeding capacity.

When analysing the seeding kinetics, the different disease durations of our selected cases must be noted. In particular, our PDD cases had longer disease duration than PD, although both shared the same Braak stage 6. The Braak staging, however, only represents the distribution of LB pathology and fails to consider the load of LBs. We have shown that PDD brains showed higher LB load in the entorhinal cortex compared to PD brains. Thus, a higher accumulation of fibrils in PDD brains might have resulted in faster seeding kinetics. Nevertheless, our DLB cases had shorter disease duration than PDD, but the comparable burden of LBs and both diseases demonstrated similar seeding kinetics. Thus, the differential seeding kinetics is likely to result from strain properties or burden of pathology rather than disease duration.

Previous studies of *α*Syn SAA with CSF showed that PD and MSA could be distinguished using *T*_50_, *V*_MAX_ (***Poggiolini et al., 2021***), and *F*_MAX_ (***Shahnawaz et al., 2020***, ***2017***). However, in our *α*Syn SAA with brain-derived fibrils, the same parameters cannot distinguish between PD and MSA. Differences in the findings are likely to arise from the type of seeds. An unprocessed CSF would likely comprise distinct and different concentrations of *α*Syn species. Moreover, the choice of biosample (CSF, olfactory mucosa, skin) significantly affects the outcome of seeding kinetics (***Chahine et al., 2020***). By isolating the brain-derived *α*Syn fibrils, we could study the seeding kinetics of the fibrils themselves and, as a result, better understand their distinct intrinsic seeding properties.

Interestingly, one healthy control case (HC 2) showed amplification towards the end of the SAA reaction. The subject had Braak *α*Syn stage 0, suggesting the absence of incidental Lewy bodies. Previously, positive seeding activities have been detected in HCs (***Han et al., 2020***). The exact cause of this observation is unclear, but the long reaction time might have contributed to the aggregation. SAA studies have shown that their HCs remain negative at 48-60 hrs (***Groveman et al., 2018***; ***Bargar et al., 2021***). Our HCs were negative at these times and only started to aggregate from 80 hrs.

Next, examining the phosphorylated brain-derived *α*Syn fibrils (p*α*Syn), we demonstrate a possible correlation between the amount of p*α*Syn and rapid seeding kinetics. Studies have shown that p*α*Syn enhances *α*Syn aggregation in vitro (SH-SY5Y cells) and in vivo (rodent models) (***Smith et al., 2005***; ***Karampetsou et al., 2017***). However, contradicting findings have also been reported, suggesting a neuroprotective role of p*α*Syn (***Ma et al., 2016***; ***Ghanem et al., 2022***). Although the biological role of p*α*Syn is unclear, we report that a higher amount of p*α*Syn may contribute to the more aggressive seeding kinetics observed in PDD and DLB brain-derived *α*Syn fibrils than those of PD and MSA.

Further evidence of ‘strain-like’ property was exhibited by the distinct biochemical profiles of the brain-derived *α*Syn fibrils from different synucleinopathies. The brain-derived *α*Syn fibrils from PD and MSA were more susceptible to GndHCl denaturation than PDD and DLB, indicating conformational instability of PD and MSA brain-derived *α*Syn fibrils. Nevertheless, it must be noted that the rapid denaturation of PD fibrils might be due to a low concentration of *α*Syn fibrils, as demonstrated by a lower Lewy pathology load in the entorhinal cortex. A previous study also showed that human brain-derived *α*Syn fibrils from DLB were more stable than MSA by using GdnHCl treatment (***Lau et al., 2020***). Interestingly, the less stable MSA fibrils propagated faster in TgM83 mice than the more stable DLB fibrils. Such inverse correlation between conformational stability and disease propagation has been characterized in prion disease (***Legname et al., 2006***). Therefore, our findings on the differential conformational stability could reflect the strain-specific characteristics of *α*Syn fibrils. Subsequently, it could be related to the fast disease progression of MSA compared to PDD and DLB.

Subsequently, by assessing the PK-resistance of the *α*Syn fibrils, we illustrate further the molecular diversity of *α*Syn fibrils. The sizes of the low MW PK-resistant fragments differed between diseases and cases within each disease. These distinct PK-resistant peptides suggest the presence of conformational variants indicative of *α*Syn fibrillar strains within a single disease entity. Similarly, Guo and colleagues reported evidence of distinct *α*Syn fibrillar conformers in different PDD patients’ brains (***Guo et al., 2013***). Our group has also shown different SAA kinetics in the CSF samples of PD patients divided into four different subtypes based on their baseline and progressive motor and non-motor symptoms (***Lawton et al., 2018***), suggesting *α*Syn strain heterogeneity within the same disease (***Poggiolini et al., 2021***). Finally, unlike PDD and DLB, MSA *α*Syn fibrils gradually disappeared upon PK treatment without being digested into small fragments. The lack of PK-resistant peptides in MSA indicates a readily digestible property of MSA *α*Syn fibrils. Together with its conformational instability, these biochemical properties might indicate a specific *α*Syn strain in MSA.

Interestingly, MSA fibrils also showed distinct structural features compared to those from PD, PDD, and DLB. The fibrils from MSA were predominantly ‘straight’, and those from PD, PDD, and DLB were ‘straight’ and ‘twisted’. While the ‘straight’ brain-derived fibrils were densely decorated with MJFR-14 and anti-p*α*Syn antibodies, confirming their identity as a phosphorylated *α*Syn fibril, the ‘twisted’ fibrils were not recognised by these proteins. Similarly, Spillantini and colleagues also reported a finding of ‘twisted’ fibril from the cingulate cortex of a DLB brain with no labelling by the *α*Syn-specific antibody (***Spillantini et al., 1998***). Further ultrastructure analysis is essential to reveal the identity of the unlabeled twisted fibrils.

The methodological limitations of the structural characterisation in this study should be considered. Cryo-electron microscope (EM) ultrastructure revealed the twisted structures of the MSA brain-derived *α*Syn fibrils and single twisted protofilament structures in PD, PDD and DLB brains (***Yang et al., 2022***). Our TEM structures may differ due to the selection of different brain regions compared to those used in the literature and the limitation of the resolving power of TEM. Future cryo-EM structural studies of *α*Syn fibrils from not only different synucleinopathies but also different brain regions and subtypes of PD are required. Accumulating evidence already suggests the presence of distinct biochemical properties among clinical and genetic subtypes of PD (***Poggiolini et al., 2021***; ***Siderowf et al., 2023***). Thus, it is essential to continue investigating the molecular profiles to understand the diversity of *α*Syn conformers and their role in pathogenesis.

### Distinct biophysical properties between brain-derived and SAA fibrils

In order to assess whether the SAA fibrils truly replicate the brain-derived fibrils (the seed), we compared the biophysical characteristics of the brain-derived *α*Syn fibrils and their respective SAA fibrils. First, by examining the resistance to GdnHCl and PK digestion, we illustrated a prominent biochemical difference between the brain-derived and SAA fibrils. High concentrations (4-5 M) of GdnHCl completely denatured the brain-derived fibrils, whilst SAA fibrils were still resistant. This indicates that the SAA fibrils are more stable and PK-resistant than their respective brain-derived fibrils. Furthermore, the SAA fibrils from all four synucleinopathies had identical patterns of PK resistance. The SAA amplifies fibrils with uniform biochemical features, losing the intrinsic properties of the original seed fibril and failing to replicate the biochemical properties of the original brain-derived fibrils.

The differences in biochemical stability might arise from the distinct distribution of the fibrils. SAA fibrils were clustered in negative-stain TEM, whereas the brain-derived fibrils were distributed in single filaments (Figure 5). Also, this study used a PIPES-based buffer to amplify the SAA fibrils (***Poggiolini et al., 2021***). As the microenvironmental context of *α*Syn amplification substantially affects the newly amplified fibrils (***Peng et al., 2018***; ***Gustavsson et al., 2021***), the artificial in vitro environment can favor the formation of specific three-dimensional structures that are more conformationally stable.

Next, we showed that brain-derived fibrils display a mixture of straight and twisted structures, while SAA fibrils have a unified morphology: straight (PD, PDD, DLB) or twisted (MSA). This was an unexpected finding, as the conformational stability patterns were identical across all SAA fibrils. Here, technical limitations exist where the conformational stability was assessed using a single antibody targeting the 91-99 region of *α*Syn. A future study could employ multiple antibodies targeting the N- and C-terminus to further understand the relations between conformational stability and fibril morphology.

The striking structural difference may be due to a dominant seeding fibril within the mixture of brain-derived fibrils. Interestingly, straight fibrils were dominant in MSA brains, but the corresponding SAA fibrils were twisted. The commonality of a particular structure in a fibrillar mixture may not be a determinant of a dominant seeding fibril. Other factors might contribute, such as metal ions (***Uversky et al., 2001***) and post-translational modifications (PTMs), which also affect seeding efficiency (***Balana et al., 2023***). Also, our PIPES-based SAA reaction buffer with 150 mM NaCl is likely to have influenced the SAA fibril structure. However, the discrete structure of the MSA SAA fibrils compared to those of PD, PDD, and DLB implies that a complex matrix of intrinsic factors within the brain-derived fraction impacts seeding amplification and the resulting fibrillar structure.

A recent cryo-EM analysis of the MSA brain-derived and SAA fibrils revealed striking structural differences (***Schweighauser et al., 2020***; ***Lövestam et al., 2021***). A few significant differences included the protofilament fold, the inter-protofilament interface and the geometry of specific residues that shape the hydrophobic core. As the TEM has restricted resolution, future cryo-EM analyses are imperative to connect the structure to the disease heterogeneity and progression.

Lastly, we investigated the differences in the phosphorylation level between the brain-derived and SAA *α*Syn fibrils to identify differences in the PTM patterns. The brain-derived fibrils generally showed a higher phosphorylation level than the SAA fibrils. PDD and DLB SAA fibrils were weakly phosphorylated and decorated with p*α*Syn immunogold labels at the rare end of the fibrils (Figure 6). Combining these findings, we can argue the possibility that the SAA fibrils extended from p*α*Syn species dissociated from the brain-derived fibrils. Therefore, the amount of initial p*α*Syn fibrils in the seed might be a critical determinant of the phosphorylation state of the resulting SAA fibrils.

The dissimilarity observed in the brain-derived and SAA fibrils might result from the RT-QuIC methodology. For instance, prion studies reported that RT-QuIC end-products are non-infectious (***Coysh and Mead, 2022***). On the other hand, PMCA prion end-products are infectious in vitro and in vivo and maintain strain-specific properties (***Castilla et al., 2005***). This indicates the limitation of RT-QuIC in reproducing the toxicity of the source seeds as it loses biological information. Therefore, the biophysical differences between the brain-derived and SAA fibrils might result from the limitations of the RT-QuIC methodology.

Studies have explored the pathogenicity of brain-derived *α*Syn fibrils in vitro using primary neurons, oligodendrocytes, and HEK cell lines expressing mutated *α*Syn (***Woerman et al., 2018a***; ***Peng et al., 2018***; ***Woerman et al., 2015***; ***Yamasaki et al., 2019***). Moreover, similar pathological characterization has been done in different mouse models (***Lau et al., 2020***; ***Woerman et al., 2019***, ***2018a***; ***Prusiner et al., 2015***; ***Peng et al., 2018***; ***Masuda-Suzukake et al., 2013***; ***Watts et al., 2013***; ***Bernis et al., 2015***; ***Woerman et al., 2018b***, ***2020***). Interestingly, both in vitro and in vivo, MSA brain-derived *α*Syn induced significant *α*Syn aggregation, whereas those from PD/PDD and DLB brains produced either no or low *α*Syn accumulation. Similar disease-specific patterns were also observed in SAA-amplified fibrils (***Shahnawaz et al., 2020***; ***Van der Perren et al., 2020***; ***Tanudjojo et al., 2021***). In future studies, it would be crucial to examine the brain-derived and SAA fibrils in comparison in cellular and mouse models for their pathological characteristics.

In summary, the distinct biochemical profiles and structures of the brain-derived *α*Syn fibrils from different synucleinopathies provide supporting evidence of molecular diversity of brain-derived *α*Syn fibrils among different synucleinopathies and individual patients. Moreover, the SAA fibrils failed to adopt the biochemical, structural and PTM properties of the seed brain-derived *α*Syn fibrils, which addresses a significant limitation of SAA in replicating the intrinsic biophysical properties of the seed fibrils. Therefore, our finding highlights the necessity of re-evaluating the SAA seeding mechanism and its capacity to generate disease-relevant *α*Syn fibrils used in different in vitro and in vivo models. Furthermore, it remains essential to investigate the human-derived *α*Syn fibrils in the different brain regions and in other peripheral sites to explore *α*Syn strains and how they may affect the progression of pathology. They also remain the most disease-specific fibrils used in cellular and animal models.

## Methods and Materials

### Patient Collection

Brain tissues from patients with PD and PDD were obtained from the Parkinson’s UK Brain Bank (Imperial College London, UK) in accordance with approved protocols by the London Multicentre Research Ethics Committee. Brain tissues from patients with DLB, MSA and HC subjects were collected from the Oxford Brain Bank (OBB, University of Oxford, UK) in accordance with approved protocols by the South Central - Oxford C Research Ethics Committee (ref 23/SC/0241). All participants had given prior written informed consent for the brain donation. Both brain banks comply with the requirements of the Human Tissue Act 2004 and the Codes of Practice set by the Human Tissue Authority (HTA licence numbers 12275 for the Imperial and 12217 for OBB). The clinico-pathological demographics of the patients and healthy controls are summarized in Table S1.

### Extraction of *α*Syn fibrils from the human brain

Sarkosyl insoluble *α*Syn fibril was extracted from the brains of PD (*n*=3), PDD (*n*=3), DLB (*n*=3), MSA (*n*=3) patients and healthy controls (*n*=3). The entorhinal cortex was selected for PD, PDD, and DLB, and the striatum was selected for MSA. The extraction protocol was adapted from the method presented by Schweighauser and colleagues (***Schweighauser et al., 2020***). Brain tissues (0.5 g) were homogenized in an extraction buffer of 10 mM Tris-HCl, 0.8 M NaCl, 10% sucrose and 1 mM EGTA (pH 7.5). Sarkosyl was added to a final concentration of 2% and incubated for 30 min at 37 °C. Homogenates were centrifuged at 10,000 g for 10 min at 4 °C. The supernatants were centrifuged at 100,000 g for 60 min at 4 °C. The pellet was washed, resuspended in 500 µl/g of extraction buffer, and centrifuged at 500 g for 1 min at 4 °C. The resulting supernatants were diluted 1/3 in buffer consisting of 50 mM Tris-HCl, 0.15 M NaCl, 10 % sucrose, and 0.2% sarkosyl (pH 7.5). The diluted supernatant was centrifuged at 100,000 g for 60 min at 4 °C. The resulting pellet was washed and resuspended in 250 µl/g of 30 mM Tris-HCl (pH 7.5), being the sarkosyl-insoluble fraction. The protein concentration was determined using the bicinchoninic acid assay (BCA) (Thermo Scientific). The sarkosyl insoluble fraction was aliquoted and stored at −80 °C.

### *α*Syn seeding amplification assay (SAA)

The brain-derived *α*Syn fibrils were diluted 1:1,000 in 30 mM Tris-HCl (pH 7.5), and 2 µl was added to 98 µl of the reaction mixture consisting of 100 mM PIPES (pH 7), 150 mM NaCl, 0.1 mg/ml of recombinant *α*Syn (rPeptide) and 10 µM thioflavin-T (ThT) to a final reaction volume of 100 µl. Before use, the recombinant *α*Syn was filtered through a 100 kDa molecular weight cut-off (MWCO) filter. The reaction mixture was loaded onto a black 96-well plate with a clear bottom (Nalgene Nunc). The plate was sealed with a plate sealing film (Thermo Scientific) and incubated at 42 °C in a BMG FLUOstar Omega plate reader for 100 h with cycles of 1 min shaking (400 rpm double orbital) and 1 min rest. ThT fluorescence measurement (450 nm excitation and 480 nm emission) was taken every 30 min. Each sample was run in six replicates. The reaction end-products were ultracentrifuged at 100,000 g for 1 h at 4 °C and collected as SAA fibril.

### Conformational stability assay and immunoblotting

The brain-derived *α*Syn fibrils (10 µg) and SAA fibrils (1 µg) were treated with different GdnHCl concentrations (0-5 M) at 37 °C for 1 h on a thermoshaker (800 rpm). The reactions were stopped by reducing the GdnHCl concentration to 0.5 M. The fibrils were treated with 1 µg/ml of proteinase-K (PK) at 37 °C for 30 min on a thermoshaker (500 rpm). Digested samples were collected using ultra-centrifugation at 100,000 g, 4 °C for 1 h. Tricine sample buffer (Bio-Rad) was added to the samples and boiled at 95 °C for 10 min. The samples were analyzed with 16.5% Tris-Tricine gels (Bio-Rad) and immunoblotted on nitrocellulose membranes (Amersham). Membranes were blocked with 5% skimmed milk in TBS-Tween and incubated with anti-*α*Syn clone 42 (BD Biosciences, 1:1,000 dilution). In a slot blot, 2 µg of protein was immobilized on a nitrocellulose membrane (Amersham) by filtration using a slot blot apparatus (GE Healthcare). The membrane was blocked with 5% skimmed milk and probed with MJFR-14-6-4-2 (Abcam, 1:3000 dilution) and EP1536Y (Abcam, 1:1,000 dilution). The membranes were developed using an ECL Western blot detection kit (Amersham). Semi-quantitative analysis of the slot blot data was performed using ImageJ.

### Transmission electron microscopy (TEM) and immunolabeling electron microscopy

Five µl of 0.5 µg brain-derived *α*Syn fibrils and 5 µl of SAA fibrils were applied to glow-discharged carbon grids (Agar Scientific, 300 mesh) and incubated for 2 min. The grid was washed with water for 10 sec and negatively stained with 2% uranyl acetate for 10 sec. Stained samples were imaged on an FEI Tecnai T12 microscope operated at 120 kV.

For immunolabeling, the fibrils were applied to glow-discharged carbon grids and blocked with a blocking buffer (0.2% fish gelatin in PBS). Then, the grid was incubated with antibodies targeting the phosphorylated *α*Syn (EP1536Y, Abcam, 1:20 dilution) and *α*Syn fibril conformer (MJFR-14-6-4-2, Abcam, 1:50 dilution). After washing with blocking buffer, the sample was incubated with goat anti-rabbit IgG coupled with 10 nm gold (Ab27234, Abcam) diluted 1:10 in the blocking buffer. Then, the grid was fixed with 0.1% glutaraldehyde in PBS. The samples were stained with 2% uranyl acetate for 30 s.

### Immunohistochemistry

Six µm-thick sections of formalin-fixed paraffin-embedded (FFPE) human brain tissue sections were de-paraffinized in xylene (3 x 5 min) and rehydrated through decreasing concentration of industrial denatured alcohol (IDA) (100%, 100%, 90%, 70%; 5 min each) and subsequently in distilled water (5 min). For antigen retrieval, sections were treated with 70% formic acid for 15 min. The sections were then blocked with 3% H_2_O_2_ for 20 min and with 10% fetal bovine serum (FBS) in PBS for 30 min. Sections were incubated with anti-*α*Syn clone 42 (BD Biosciences) diluted 1:1,000 in 10% FBS overnight at 4 °C. After washing with PBS (3 x 5 min), sections were incubated with Ms/Rb-HRP secondary antibody (Agilent Technologies) for 30 min at RT. Sections were developed with Dako DAB (Agilent Technologies) and counterstained with haematoxylin. Finally, sections were mounted to coverslips with DPX (Thermo Scientific).

### Statistical analysis

Mann-Whitney U test was used when comparing two independent groups. Independent Kruskal-Wallis Test was performed to compare more than three groups for SAA kinetic parameters Figure 2, fibril dimensions (PD, PDD, DLB, and MSA) Figure 5, and the amount of phosphorylated *α*Syn fibrils Figure 6. *P* values < 0.05 were considered statistically significant. All statistical analyses were performed using SPSS Statistics, Version 28.

## Acknowledgments

We thank all the patients and their families for the valuable brain donations for research. We also want to thank Dr. Errin Johnson and the Oxford bioimaging facility for providing access and training for transmission electron microscopy and assisting with acquiring the imaging data. We acknowledge Dr. Alan King Lun Liu for helping us with the case selection and analysis of the clinical summary of the patients. We acknowledge the Oxford Brain Bank, supported by Brains for Dementia Research (BDR) (Alzheimer Society and Alzheimer Research UK) and the National Institute for Health Research (NIHR) Oxford Biomedical Research Centre (BRC). LP is supported by the GSK-Institute of Molecular & Computational Medicine, NIHR Oxford BRC, Parkinson Foundation, the Michael J. Fox Foundation, the Galen and Hilary Weston Foundation, and the National Institute of Health. LC is supported by NIHR Oxford BRC.

## Supplementary Material

### Cases

The clinical diagnosis of Parkinson’s disease patients was performed using the UK Parkinson’s Disease Brain Bank criteria, followed by a neurologist review (***Daniel and Lees, 1993***). PDD patients were diagnosed using the Movement Disorder Society task force PDD diagnostic criteria (***Dubois et al., 2007***). DLB patients were diagnosed using the DLB consortium diagnostic criteria (***McKeith et al., 2017***). MSA patients were diagnosed based on the MSA consensus statement (***Gilman et al., 1998***). All PD, PDD and DLB subjects had neuropathologically confirmed Lewy body pathology. Neuropathological diagnosis of MSA subjects was based on degeneration of striatonigral and olivopontocerebellar regions combined with glial cytoplasmic aSyn inclusions (***Papp et al., 1989***). Braak *α*Syn and tau stages were assessed according to the protocol outlined by BrainNet Europe (***Alafuzoff et al., 2008***, ***2009***). Specifically, Braak *α*Syn stages were assigned by examining the distribution of *α*Syn inclusions in the medulla, pons, midbrain, basal ganglia, hippocampus, cingulate gyrus, temporal, frontal and parietal cortices. Braak tau stages were assigned based on tau immunostains across regions of the visual cortex, the middle temporal gyrus, the anterior hippocampus and the posterior hippocampus.

**Table S1.**
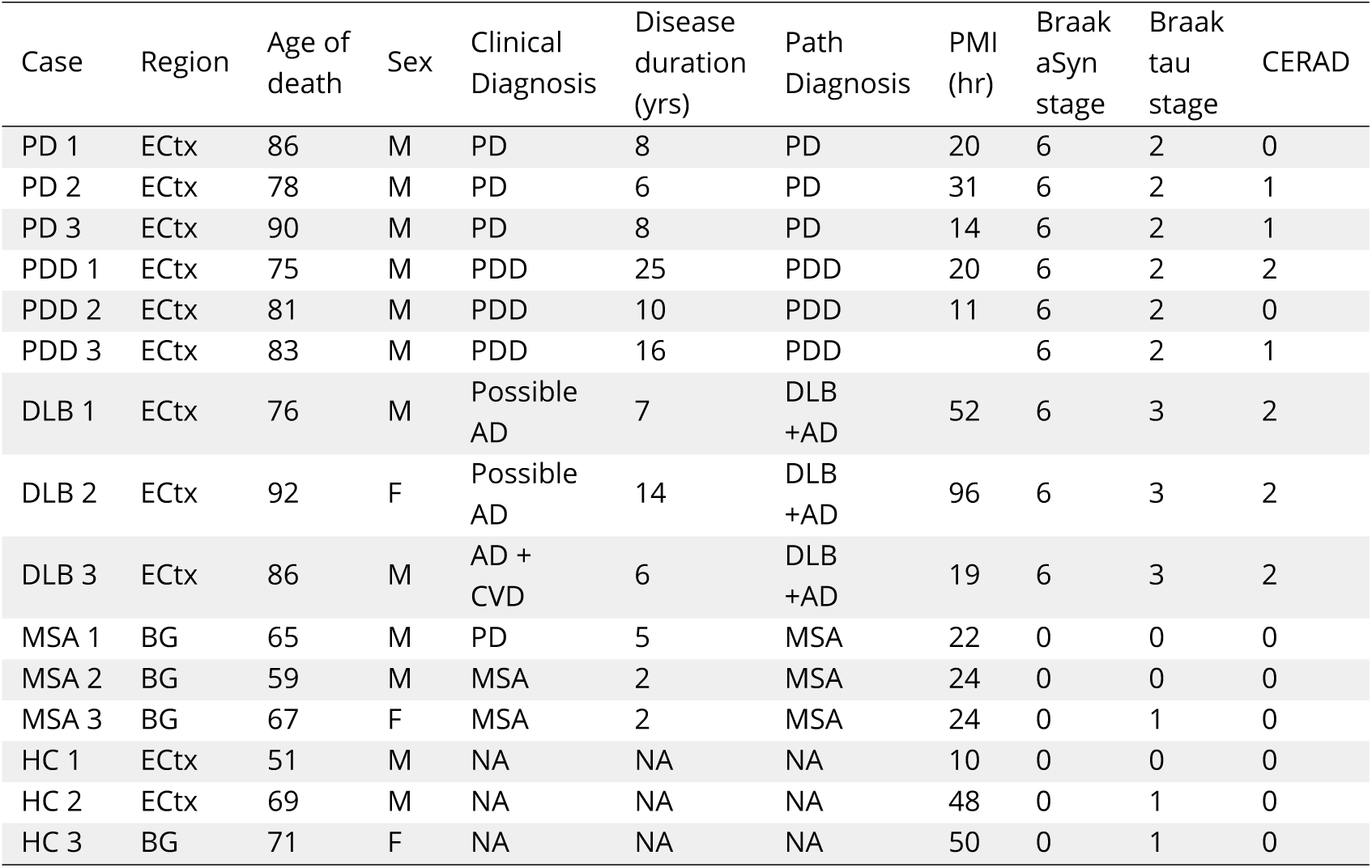
Clinicopathological summary of the patients included in this study. The table summarizes the clinical and pathological reports of the patients included in this study. PD, Parkinson’s disease; PDD, Parkinson’s disease with dementia; DLB, dementia with Lewy body; MSA, multiple system atrophy; HC, healthy control; ECtx, entorhinal cortex; BG, basal ganglia; M, male; F, female; AD, Alzheimer’s disease; CVD, cardiovascular disease; *α*Syn, *α*-synuclein.

| Case | Region | Age of death | Sex | Clinical Diagnosis | Disease duration (yrs) | Path Diagnosis | PMI (hr) | Braak αSyn stage | Braak tau stage | CERAD |
| --- | --- | --- | --- | --- | --- | --- | --- | --- | --- | --- |
| PD 1 | ECtx | 86 | M | PD | 8 | PD | 20 | 6 | 2 | 0 |
| PD 2 | ECtx | 78 | M | PD | 6 | PD | 31 | 6 | 2 | 1 |
| PD 3 | ECtx | 90 | M | PD | 8 | PD | 14 | 6 | 2 | 1 |
| PDD 1 | ECtx | 75 | M | PDD | 25 | PDD | 20 | 6 | 2 | 2 |
| PDD 2 | ECtx | 81 | M | PDD | 10 | PDD | 11 | 6 | 2 | 0 |
| PDD 3 | ECtx | 83 | M | PDD | 16 | PDD |  | 6 | 2 | 1 |
| DLB 1 | ECtx | 76 | M | Possible AD | 7 | DLB +AD | 52 | 6 | 3 | 2 |
| DLB 2 | ECtx | 92 | F | Possible AD | 14 | DLB +AD | 96 | 6 | 3 | 2 |
| DLB 3 | ECtx | 86 | M | AD + CVD | 6 | DLB +AD | 19 | 6 | 3 | 2 |
| MSA 1 | BG | 65 | M | PD | 5 | MSA | 22 | 0 | 0 | 0 |
| MSA 2 | BG | 59 | M | MSA | 2 | MSA | 24 | 0 | 0 | 0 |
| MSA 3 | BG | 67 | F | MSA | 2 | MSA | 24 | 0 | 1 | 0 |
| HC 1 | ECtx | 51 | M | NA | NA | NA | 10 | 0 | 0 | 0 |
| HC 2 | ECtx | 69 | M | NA | NA | NA | 48 | 0 | 1 | 0 |
| HC 3 | BG | 71 | F | NA | NA | NA | 50 | 0 | 1 | 0 |

**Figure S1.**
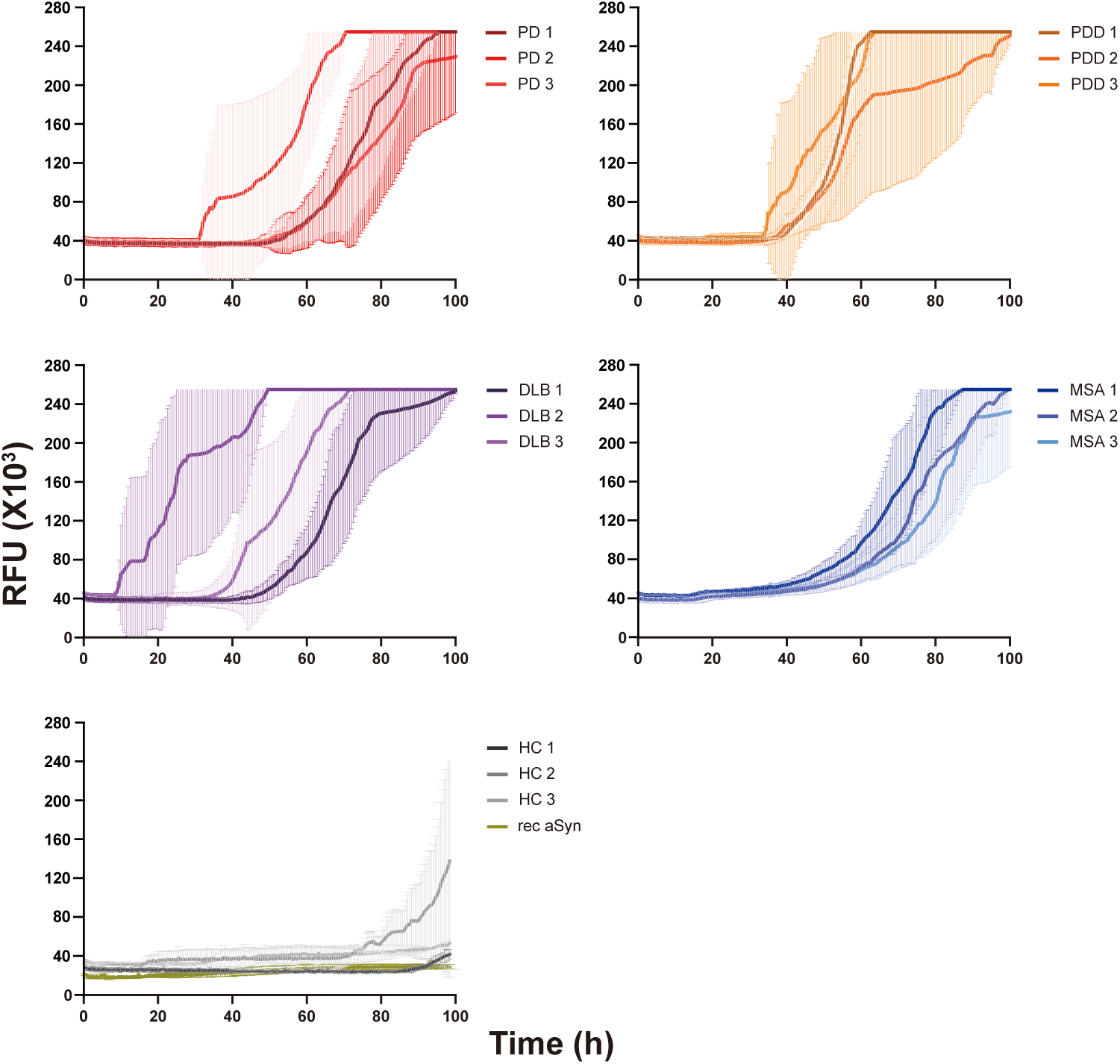
Raw data of the *α*Syn SAA reaction seeded with *α*Syn fibrils from PD, PDD, DLB, MSA, and healthy control brains. The curves represent the average of 6 replicates, and error bars indicate ± one standard deviation (SD). RFU, relative fluorescence unit.

**Figure S2.**
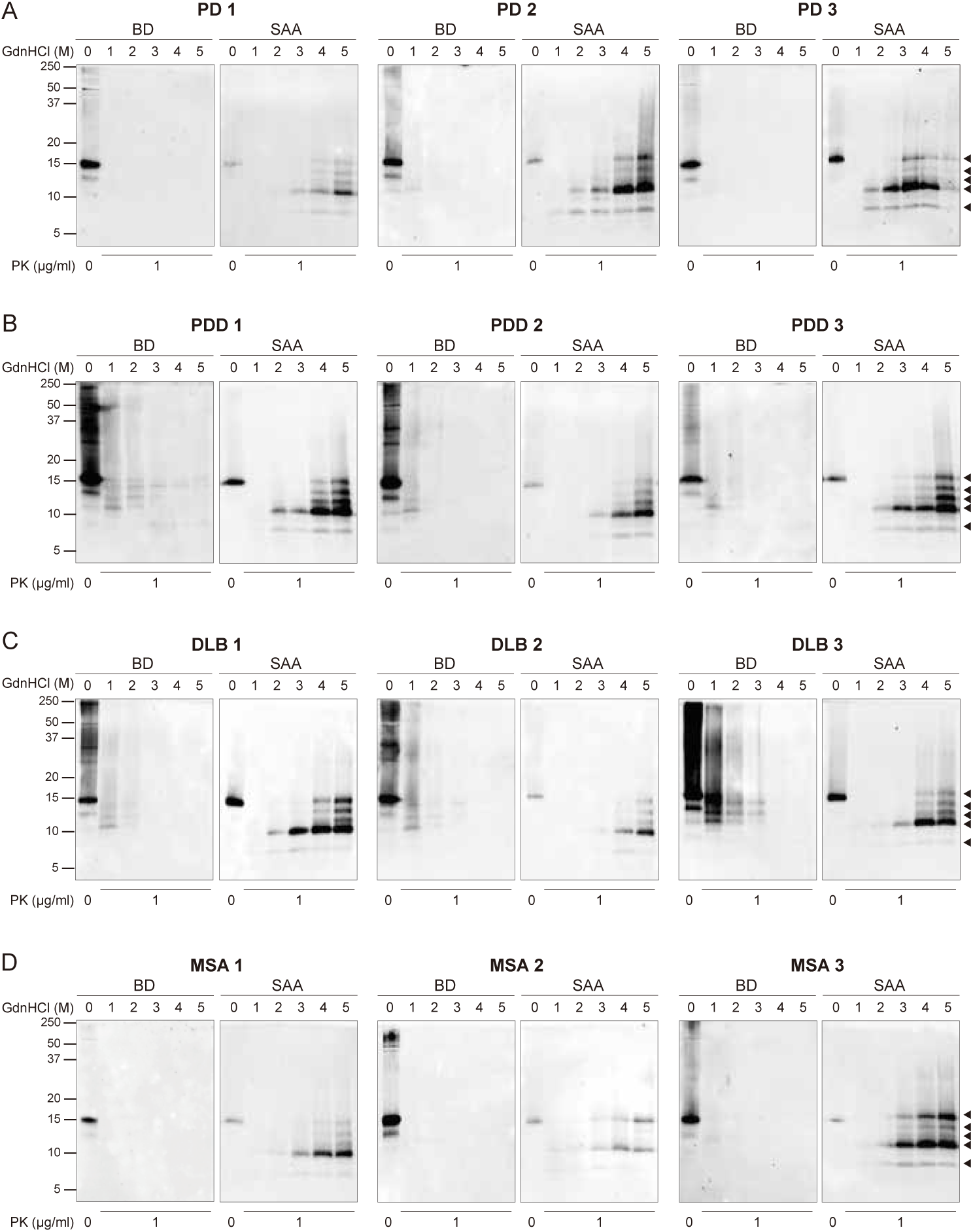
The full biochemical profiles of the brain-derived and SAA *α*Syn fibrils from all the cases. The brain-derived (BD) and SAA *α*Syn fibrils from all 3 cases of (**A**) PD, (**B**) PDD, (**C**) DLB, and (**D**) MSA were treated with GdnHCl (0-5 M) and PK (1 µ g/ml). The antibody clone 42 (BD Biosciences) was used to reveal the PK-resistant peptides. Black arrows mark the five PK-resistant peptides revealed in the SAA fibrils. The molecular weights of the protein standards are shown in kilodaltons (kDa). SAA, seeding amplification assay; GndHCl, guanidine-hydrochloride; PK, proteinase-K.

**Figure S3.**
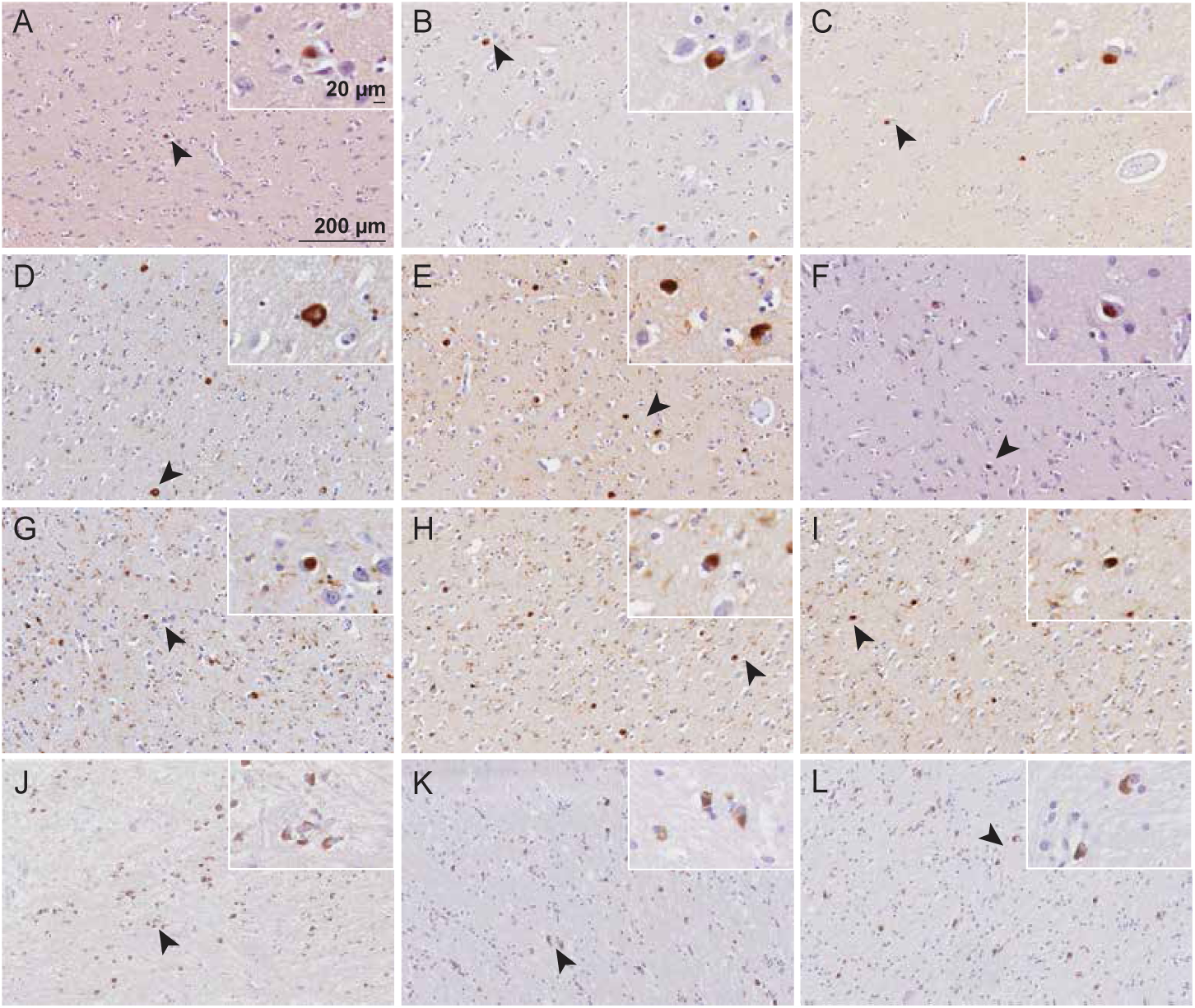
Immunohistochemical detection of *α*Syn in PD, PDD, DLB, and MSA brains. The entorhinal cortex (PD, PDD, DLB) and striatum (MSA) were stained with total *α*Syn using antibody clone 42 (BD Biosciences). (**A-C**) PD cases 1-3. (**D-F**) PDD cases 1-3. (**G-I**) DLB cases 1-3. (**J-L**) MSA cases 1-3. Arrows indicate magnified areas. Scale bars = 20 µm and 200 µm.

**Figure S4.**
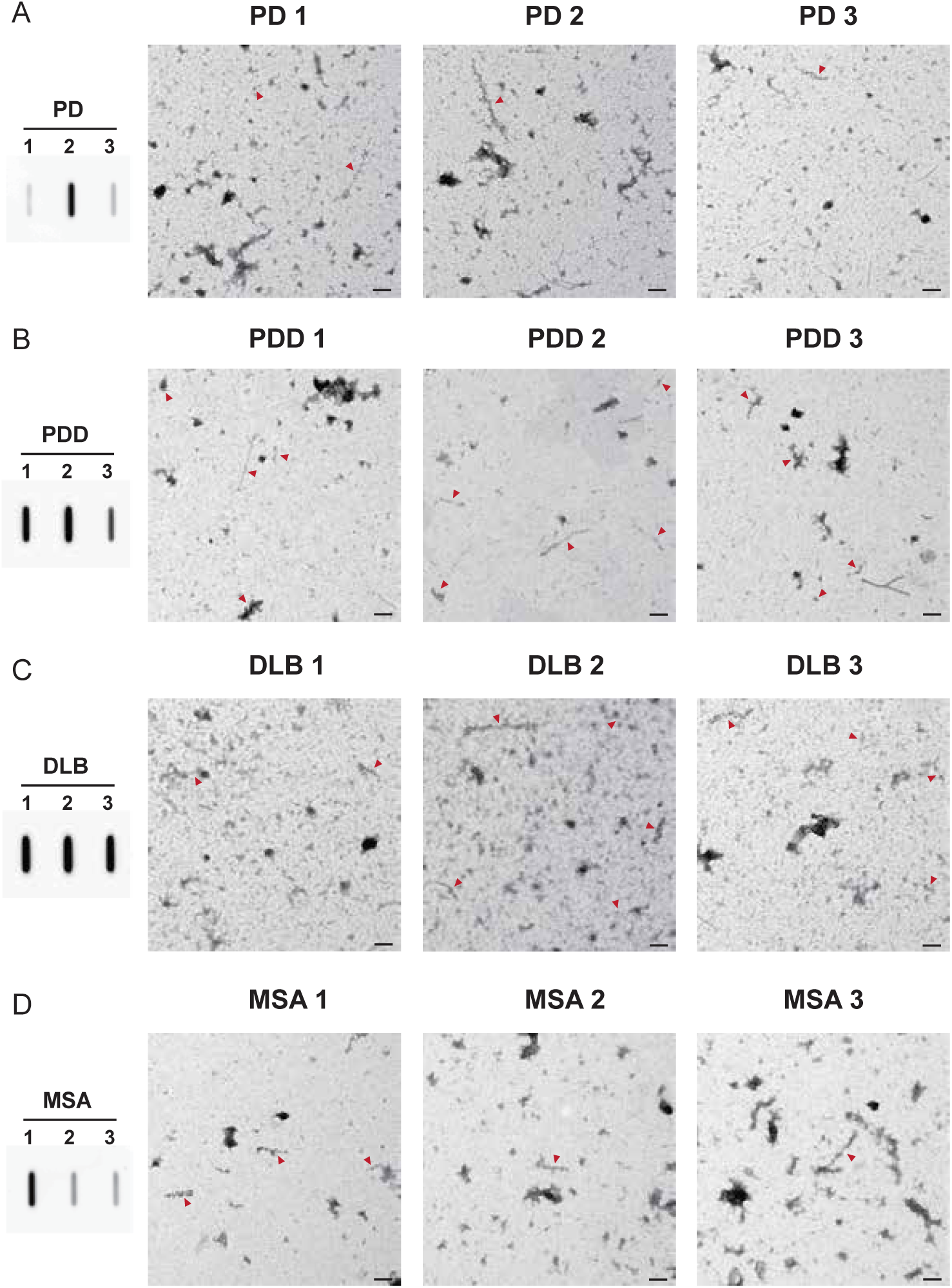
pS129 immunogold TEM images of the brain-derived *α*Syn fibrils. The brain-derived *α*Syn fibrils were labeled with anti-p*α*Syn (pS129) and imaged at low magnification to compare the relative quantities of the p*α*Syn fibrils between the diseases. All three cases were imaged for each disease (**A**) PD, (**B**) PDD, (**C**) DLB, and (**D**) MSA. The amount of p*α*Syn fibrils was examined using slot blots. Red arrows indicate labeled fibrils. Scale bar = 200 µm.

**Figure S5.**
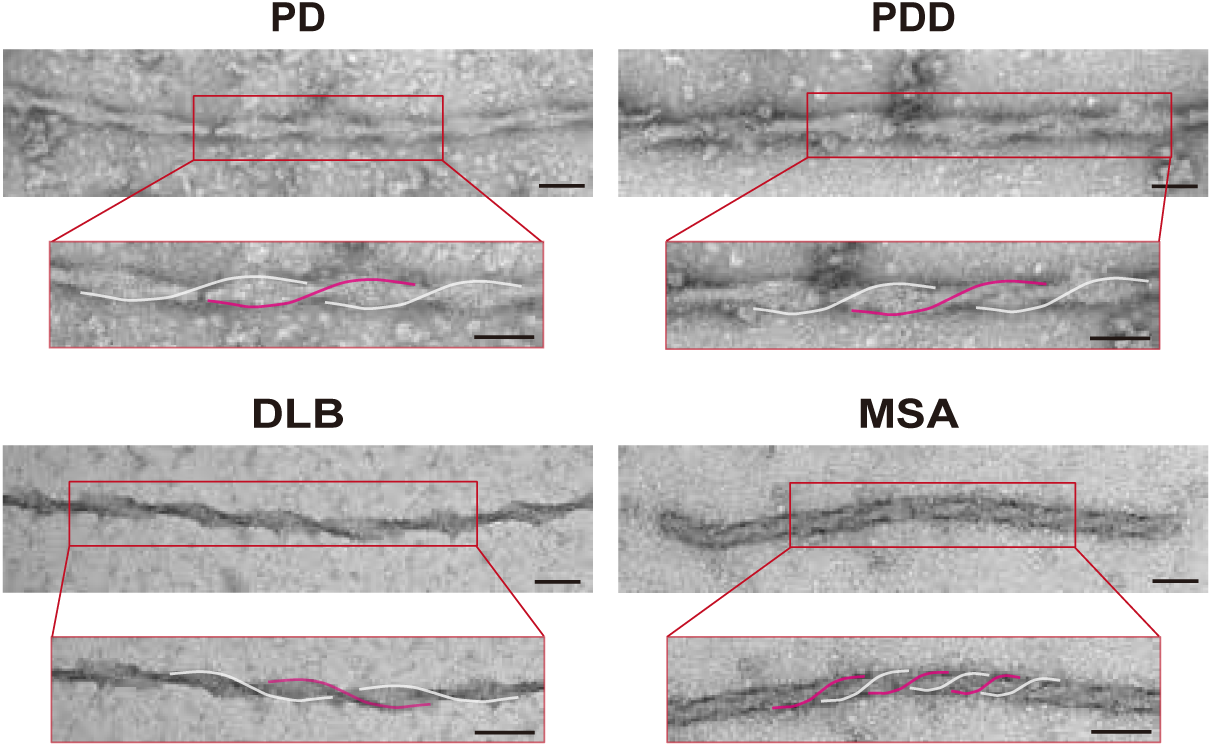
TEM images of the twisted brain-derived *α*Syn fibrils. PD, PDD, DLB, and MSA brain-derived fibrils displayed intertwined fibrils with twists. Helical twists are highlighted in pink and white. Scale bar = 50 µm.

**Figure S6.**
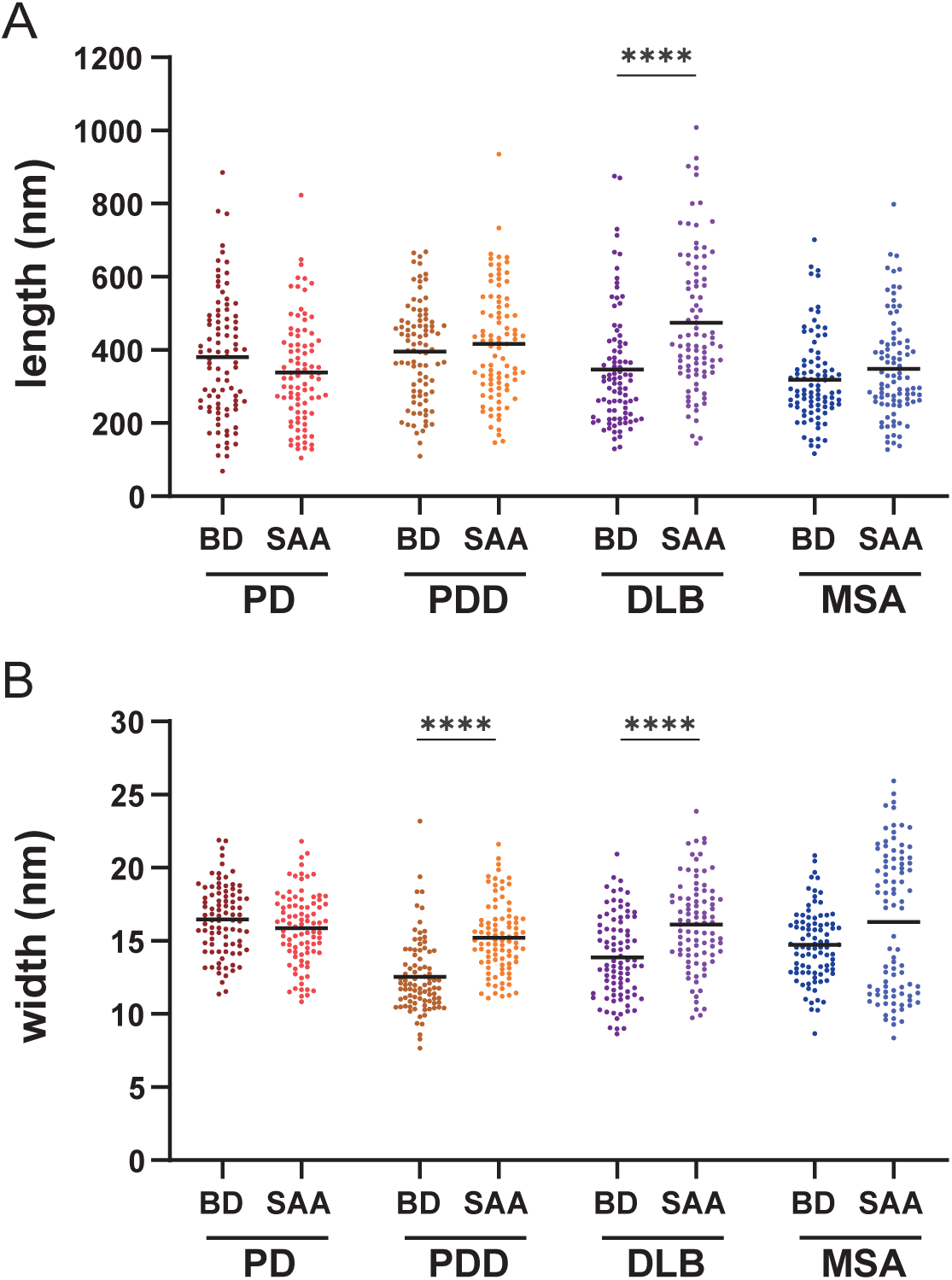
Comparison of the length and width of the brain-derived and SAA *α*Syn fibrils. Comparison of the (**A**) lengths and the (**B**) widths of the brain-derived and SAA *α*Syn fibrils. Thirty fibrils were counted for each case. Thus, a total of 90 data points were plotted (*N* = 90). The black bold line indicates an average. BD, brain-derived; SAA, seeding amplification assay. **** *P* ≤ 0.001.

## References

Alafuzoff, I., Arzberger, T., Al-Sarraj, S., Bodi, I., Bogdanovic, N., Braak, H., Bugiani, O., Del-Tredici, K., Ferrer, I., Gelpi, E., et al. (2008). Staging of neurofibrillary pathology in alzheimer’s disease: a study of the brainnet europe consortium. Brain pathology, 18(4):484–496.

Alafuzoff, I., Ince, P. G., Arzberger, T., Al-Sarraj, S., Bell, J., Bodi, I., Bogdanovic, N., Bugiani, O., Ferrer, I., Gelpi, E., et al. (2009). Staging/typing of lewy body related *α*-synuclein pathology: a study of the brainnet europe consortium. Acta neuropathologica, 117:635–652.

Altay, M. F., Liu, A. K. L., Holton, J. L., Parkkinen, L., and Lashuel, H. A. (2022). Prominent astrocytic alpha-synuclein pathology with unique post-translational modification signatures unveiled across lewy body disorders. Acta Neuropathologica Communications, 10(1):163.

Ayers, J. I., Lee, J., Monteiro, O., Woerman, A. L., Lazar, A. A., Condello, C., Paras, N. A., and Prusiner, S. B. (2022). Different α-synuclein prion strains cause dementia with lewy bodies and multiple system atrophy. Proc Natl Acad Sci U S A, 119(6).

Balana, A. T., Mahul-Mellier, A.-L., Nguyen, B. A., Horvath, M., Javed, A., Hard, E. R., Jasiqi, Y., Singh, P., Afrin, S., Pedretti, R., et al. (2023). O-glcnac modification forces the formation of an *α*-synuclein amyloid-strain with notably diminished seeding activity and pathology. bioRxiv, pages 2023–03.

Bargar, C., Wang, W., Gunzler, S. A., LeFevre, A., Wang, Z., Lerner, A. J., Singh, N., Tatsuoka, C., Appleby, B., Zhu, X., et al. (2021). Streamlined alpha-synuclein rt-quic assay for various biospecimens in parkinson’s disease and dementia with lewy bodies. Acta neuropathologica communications, 9:1–13.

Bernis, M. E., Babila, J. T., Breid, S., Wusten, K. A., Wullner, U., and Tamguney, G. (2015). Prion-like propagation of human brain-derived alpha-synuclein in transgenic mice expressing human wild-type alpha-synuclein. Acta Neuropathol Commun, 3:75.

Bousset, L., Pieri, L., Ruiz-Arlandis, G., Gath, J., Jensen, P. H., Habenstein, B., Madiona, K., Olieric, V., Bockmann, A., Meier, B. H., and Melki, R. (2013). Structural and functional characterization of two alpha-synuclein strains. Nat Commun, 4:2575.

Braak, H., Del Tredici, K., Rub, U., de Vos, R. A., Jansen Steur, E. N., and Braak, E. (2003). Staging of brain pathology related to sporadic parkinson’s disease. Neurobiol Aging, 24(2):197–211.

Castilla, J., Saá, P., Hetz, C., and Soto, C. (2005). In vitro generation of infectious scrapie prions. Cell, 121(2):195– 206.

Chahine, L. M., Beach, T. G., Brumm, M. C., Adler, C. H., Coffey, C. S., Mosovsky, S., Caspell-Garcia, C., Serrano, G. E., Munoz, D. G., III, C. L. W., Crary, J. F., Jennings, D., Taylor, P., Foroud, T., Arnedo, V., Kopil, C. M., Riley, L., Dave, K. D., Mollenhauer, B., and for the Systemic Synuclein Sampling Study (2020). In vivo distribution of α-synuclein in multiple tissues and biofluids in parkinson disease. Neurology, 95(9):e1267–e1284.

Concha-Marambio, L., Pritzkow, S., Shahnawaz, M., Farris, C. M., and Soto, C. (2023). Seed amplification assay for the detection of pathologic alpha-synuclein aggregates in cerebrospinal fluid. Nature protocols, 18(4):1179– 1196.

Coysh, T. and Mead, S. (2022). The future of seed amplification assays and clinical trials. Frontiers in Aging Neuroscience, 14:872629.

Daniel, S. and Lees, A. (1993). Parkinson’s disease society brain bank, london: overview and research. Journal of neural transmission. Supplementum, 39:165–172.

Dubois, B., Burn, D., Goetz, C., Aarsland, D., Brown, R. G., Broe, G. A., Dickson, D., Duyckaerts, C., Cummings, J., Gauthier, S., et al. (2007). Diagnostic procedures for parkinson’s disease dementia: recommendations from the movement disorder society task force. Movement disorders, 22(16):2314–2324.

Fairfoul, G., McGuire, L. I., Pal, S., Ironside, J. W., Neumann, J., Christie, S., Joachim, C., Esiri, M., Evetts, S. G., Rolinski, M., Baig, F., Ruffmann, C., Wade-Martins, R., Hu, M. T., Parkkinen, L., and Green, A. J. (2016). Alpha-synuclein rt-quic in the csf of patients with alpha-synucleinopathies. Ann Clin Transl Neurol, 3(10):812–818.

Forno, L. S. (1988). The neuropathology of parkinson’s disease. Progress in Parkinson Research, pages 11–21.

Frieg, B., Geraets, J. A., Strohäker, T., Dienemann, C., Mavroeidi, P., Jung, B. C., Kim, W. S., Lee, S.-J., Xilouri, M., Zweckstetter, M., and Schröder, G. F. (2022). Quaternary structure of patient-homogenate amplified α-synuclein fibrils modulates seeding of endogenous α-synuclein. Communications Biology, 5(1):1040.

Ghanem, S. S., Majbour, N. K., Vaikath, N. N., Ardah, M. T., Erskine, D., Jensen, N. M., Fayyad, M., Sudhakaran, I. P., Vasili, E., Melachroinou, K., et al. (2022). *α*-synuclein phosphorylation at serine 129 occurs after initial protein deposition and inhibits seeded fibril formation and toxicity. Proceedings of the National Academy of Sciences, 119(15):e2109617119.

Gilman, S., Low, P., Quinn, N., Albanese, A., Ben-Shlomo, Y., Fowler, C., Kaufmann, H., Klockgether, T., Lang, A., Lantos, P., et al. (1998). Consensus statement on the diagnosis of multiple system atrophy. Clinical autonomic research, 8:359–362.

Groveman, B. R., Orru, C. D., Hughson, A. G., Raymond, L. D., Zanusso, G., Ghetti, B., Campbell, K. J., Safar, J., Galasko, D., and Caughey, B. (2018). Rapid and ultra-sensitive quantitation of disease-associated alpha-synuclein seeds in brain and cerebrospinal fluid by alphasyn rt-quic. Acta Neuropathol Commun, 6(1):7.

Guerrero-Ferreira, R., Taylor, N. M., Arteni, A. A., Kumari, P., Mona, D., Ringler, P., Britschgi, M., Lauer, M. E., Makky, A., Verasdonck, J., Riek, R., Melki, R., Meier, B. H., Bockmann, A., Bousset, L., and Stahlberg, H. (2019). Two new polymorphic structures of human full-length alpha-synuclein fibrils solved by cryo-electron microscopy. Elife, 8.

Guo, J. L., Covell, D. J., Daniels, J. P., Iba, M., Stieber, A., Zhang, B., Riddle, D. M., Kwong, L. K., Xu, Y., Trojanowski, J. Q., and Lee, V. M. (2013). Distinct α-synuclein strains differentially promote tau inclusions in neurons. Cell, 154(1):103–17.

Gustavsson, N., Savchenko, E., Klementieva, O., and Roybon, L. (2021). The intracellular milieu of Parkinson’s disease patient brain cells modulates alpha-synuclein protein aggregation. Acta Neuropathologica Communications, 9(1):1–6.

Han, J.-Y., Jang, H.-S., Green, A. J., and Choi, Y. P. (2020). Rt-quic-based detection of alpha-synuclein seeding activity in brains of dementia with lewy body patients and of a transgenic mouse model of synucleinopathy. Prion, 14(1):88–94.

Holec, S. A. M., Lee, J., Oehler, A., Ooi, F. K., Mordes, D. A., Olson, S. H., Prusiner, S. B., and Woerman, A. L. (2022). Multiple system atrophy prions transmit neurological disease to mice expressing wild-type human α-synuclein. Acta Neuropathol, 144(4):677–690.

Karampetsou, M., Ardah, M. T., Semitekolou, M., Polissidis, A., Samiotaki, M., Kalomoiri, M., Majbour, N., Xanthou, G., El-Agnaf, O. M. A., and Vekrellis, K. (2017). Phosphorylated exogenous alpha-synuclein fibrils exacerbate pathology and induce neuronal dysfunction in mice. Scientific Reports, 7(1):16533.

Kuzkina, A., Bargar, C., Schmitt, D., Rößle, J., Wang, W., Schubert, A.-L., Tatsuoka, C., Gunzler, S. A., Zou, W.-Q., Volkmann, J., Sommer, C., Doppler, K., and Chen, S. G. (2021). Diagnostic value of skin rt-quic in parkinson’s disease: a two-laboratory study. npj Parkinson’s Disease, 7(1):99.

Lau, A., So, R. W. L., Lau, H. H. C., Sang, J. C., Ruiz-Riquelme, A., Fleck, S. C., Stuart, E., Menon, S., Visanji, N. P., Meisl, G., Faidi, R., Marano, M. M., Schmitt-Ulms, C., Wang, Z., Fraser, P. E., Tandon, A., Hyman, B. T., Wille, H., Ingelsson, M., Klenerman, D., and Watts, J. C. (2020). alpha-synuclein strains target distinct brain regions and cell types. Nat Neurosci, 23(1):21–31.

Lawton, M., Ben-Shlomo, Y., May, M. T., Baig, F., Barber, T. R., Klein, J. C., Swallow, D. M., Malek, N., Grosset, K. A., and Bajaj, N. (2018). Developing and validating parkinson’s disease subtypes and their motor and cognitive progression. Journal of Neurology, Neurosurgery & Psychiatry, 89(12):1279–1287.

Legname, G., Nguyen, H.-O. B., Peretz, D., Cohen, F. E., DeArmond, S. J., and Prusiner, S. B. (2006). Continuum of prion protein structures enciphers a multitude of prion isolate-specified phenotypes. Proceedings of the National Academy of Sciences, 103(50):19105–19110.

Li, B., Ge, P., Murray, K. A., Sheth, P., Zhang, M., Nair, G., Sawaya, M. R., Shin, W. S., Boyer, D. R., Ye, S., Eisenberg, D. S., Zhou, Z. H., and Jiang, L. (2018). Cryo-em of full-length α-synuclein reveals fibril polymorphs with a common structural kernel. Nat Commun, 9(1):3609.

Lövestam, S., Schweighauser, M., Matsubara, T., Murayama, S., Tomita, T., Ando, T., Hasegawa, K., Yoshida, M., Tarutani, A., and Hasegawa, M. (2021). Seeded assembly in vitro does not replicate the structures of α-synuclein filaments from multiple system atrophy. FEBS Open bio, 11(4):999–1013.

Ma, M.-R., Hu, Z.-W., Zhao, Y.-F., Chen, Y.-X., and Li, Y.-M. (2016). Phosphorylation induces distinct alpha-synuclein strain formation. Scientific Reports, 6(1):37130.

Mammana, A., Baiardi, S., Rossi, M., Franceschini, A., Donadio, V., Capellari, S., Caughey, B., and Parchi, P. (2020). Detection of prions in skin punch biopsies of creutzfeldt-jakob disease patients. Ann Clin Transl Neurol, 7(4):559–564.

Masuda-Suzukake, M., Nonaka, T., Hosokawa, M., Oikawa, T., Arai, T., Akiyama, H., Mann, D. M., and Hasegawa, M. (2013). Prion-like spreading of pathological α-synuclein in brain. Brain, 136(Pt 4):1128–38.

McKeith, I. G., Boeve, B. F., Dickson, D. W., Halliday, G., Taylor, J.-P., Weintraub, D., Aarsland, D., Galvin, J., Attems, J., Ballard, C. G., et al. (2017). Diagnosis and management of dementia with lewy bodies: Fourth consensus report of the dlb consortium. Neurology, 89(1):88–100.

Melki, R. (2017). How the shapes of seeds can influence pathology. AA.

Papp, M. I., Kahn, J. E., and Lantos, P. L. (1989). Glial cytoplasmic inclusions in the cns of patients with multiple system atrophy (striatonigral degeneration, olivopontocerebellar atrophy and shy-drager syndrome). J Neurol Sci, 94(1-3):79–100.

Peelaerts, W., Bousset, L., Van der Perren, A., Moskalyuk, A., Pulizzi, R., Giugliano, M., Van den Haute, C., Melki, R., and Baekelandt, V. (2015). alpha-synuclein strains cause distinct synucleinopathies after local and systemic administration. Nature, 522(7556):340–4.

Peng, C., Gathagan, R. J., Covell, D. J., Medellin, C., Stieber, A., Robinson, J. L., Zhang, B., Pitkin, R. M., Olufemi, M. F., Luk, K. C., Trojanowski, J. Q., and Lee, V. M. (2018). Cellular milieu imparts distinct pathological alpha-synuclein strains in alpha-synucleinopathies. Nature, 557(7706):558–563.

Poggiolini, I., Gupta, V., Lawton, M., Lee, S., El-Turabi, A., Querejeta-Coma, A., Trenkwalder, C., Sixel-Döring, F., Foubert-Samier, A., Le Traon, A. P., Plazzi, G., Biscarini, F., Montplaisir, J., Gagnon, J. F., Postuma, R. B., Antelmi, E., Meissner, W. G., Mollenhauer, B., Ben-Shlomo, Y., Hu, M. T., and Parkkinen, L. (2021). Diagnostic value of cerebrospinal fluid alpha-synuclein seed quantification in synucleinopathies. Brain.

Prusiner, S. B., Woerman, A. L., Mordes, D. A., Watts, J. C., Rampersaud, R., Berry, D. B., Patel, S., Oehler, A., Lowe, J. K., Kravitz, S. N., Geschwind, D. H., Glidden, D. V., Halliday, G. M., Middleton, L. T., Gentleman, S. M., Grinberg, L. T., and Giles, K. (2015). Evidence for α-synuclein prions causing multiple system atrophy in humans with parkinsonism. Proc Natl Acad Sci U S A, 112(38):E5308–17.

Raymond, G. J., Race, B., Orrú, C. D., Raymond, L. D., Bongianni, M., Fiorini, M., Groveman, B. R., Ferrari, S., Sacchetto, L., Hughson, A. G., et al. (2020). Transmission of cjd from nasal brushings but not spinal fluid or rt-quic product. Annals of Clinical and Translational Neurology, 7(6):932–944.

Sano, K., Atarashi, R., Ishibashi, D., Nakagaki, T., Satoh, K., and Nishida, N. (2014). Conformational properties of prion strains can be transmitted to recombinant prion protein fibrils in real-time quaking-induced conversion. J Virol, 88(20):11791–801.

Sano, K., Atarashi, R., Satoh, K., Ishibashi, D., Nakagaki, T., Iwasaki, Y., Yoshida, M., Murayama, S., Mishima, K., and Nishida, N. (2018). Prion-like seeding of misfolded α-synuclein in the brains of dementia with lewy body patients in rt-quic. Mol Neurobiol, 55(5):3916–3930.

Schweighauser, M., Shi, Y., Tarutani, A., Kametani, F., Murzin, A. G., Ghetti, B., Matsubara, T., Tomita, T., Ando, T., Hasegawa, K., Murayama, S., Yoshida, M., Hasegawa, M., Scheres, S. H. W., and Goedert, M. (2020). Structures of α-synuclein filaments from multiple system atrophy. Nature.

Shahnawaz, M., Mukherjee, A., Pritzkow, S., Mendez, N., Rabadia, P., Liu, X., Hu, B., Schmeichel, A., Singer, W., Wu, G., Tsai, A. L., Shirani, H., Nilsson, K. P. R., Low, P. A., and Soto, C. (2020). Discriminating alpha-synuclein strains in parkinson’s disease and multiple system atrophy. Nature, 578(7794):273–277.

Shahnawaz, M., Tokuda, T., Waragai, M., Mendez, N., Ishii, R., Trenkwalder, C., Mollenhauer, B., and Soto, C. (2017). Development of a biochemical diagnosis of parkinson disease by detection of alpha-synuclein misfolded aggregates in cerebrospinal fluid. JAMA Neurol, 74(2):163–172.

Siderowf, A., Concha-Marambio, L., Lafontant, D.-E., Farris, C. M., Ma, Y., Urenia, P. A., Nguyen, H., Alcalay, R. N., Chahine, L. M., Foroud, T., et al. (2023). Assessment of heterogeneity among participants in the parkinson’s progression markers initiative cohort using *α*-synuclein seed amplification: a cross-sectional study. The Lancet Neurology, 22(5):407–417.

Smith, W. W., Margolis, R. L., Li, X., Troncoso, J. C., Lee, M. K., Dawson, V. L., Dawson, T. M., Iwatsubo, T., and Ross, C. A. (2005). *α*-synuclein phosphorylation enhances eosinophilic cytoplasmic inclusion formation in sh-sy5y cells. Journal of Neuroscience, 25(23):5544–5552.

Spillantini, M. G., Crowther, R. A., Jakes, R., Hasegawa, M., and Goedert, M. (1998). alpha-synuclein in filamentous inclusions of lewy bodies from parkinson’s disease and dementia with lewy bodies. Proc Natl Acad Sci U S A, 95(11):6469–73.

Spillantini, M. G., Schmidt, M. L., Lee, V. M., Trojanowski, J. Q., Jakes, R., and Goedert, M. (1997). Alpha-synuclein in lewy bodies. Nature, 388(6645):839–40.

Strohaker, T., Jung, B. C., Liou, S. H., Fernandez, C. O., Riedel, D., Becker, S., Halliday, G. M., Bennati, M., Kim, W. S., Lee, S. J., and Zweckstetter, M. (2019). Structural heterogeneity of alpha-synuclein fibrils amplified from patient brain extracts. Nat Commun, 10(1):5535.

Tanudjojo, B., Shaikh, S. S., Fenyi, A., Bousset, L., Agarwal, D., Marsh, J., Zois, C., Heman-Ackah, S., Fischer, R., Sims, D., Melki, R., and Tofaris, G. K. (2021). Phenotypic manifestation of α-synuclein strains derived from parkinson’s disease and multiple system atrophy in human dopaminergic neurons. Nature Communications, 12(1):3817.

Uversky, V. N., Li, J., and Fink, A. L. (2001). Metal-triggered structural transformations, aggregation, and fibrillation of human *α*-synuclein: a possible molecular link between parkinson’s disease and heavy metal exposure. Journal of Biological Chemistry, 276(47):44284–44296.

Van der Perren, A., Gelders, G., Fenyi, A., Bousset, L., Brito, F., Peelaerts, W., Van den Haute, C., Gentleman, S., Melki, R., and Baekelandt, V. (2020). The structural differences between patient-derived α-synuclein strains dictate characteristics of parkinson’s disease, multiple system atrophy and dementia with lewy bodies. Acta Neuropathol, 139(6):977–1000.

Watts, J. C., Giles, K., Oehler, A., Middleton, L., Dexter, D. T., Gentleman, S. M., DeArmond, S. J., and Prusiner, S. B. (2013). Transmission of multiple system atrophy prions to transgenic mice. Proc Natl Acad Sci U S A, 110(48):19555–60.

Woerman, A. L., Kazmi, S. A., Patel, S., Aoyagi, A., Oehler, A., Widjaja, K., Mordes, D. A., Olson, S. H., and Prusiner, S. B. (2018a). Familial parkinson’s point mutation abolishes multiple system atrophy prion replication. Proceedings of the National Academy of Sciences, 115(2):409–414.

Woerman, A. L., Kazmi, S. A., Patel, S., Freyman, Y., Oehler, A., Aoyagi, A., Mordes, D. A., Halliday, G. M., Middleton, L. T., Gentleman, S. M., Olson, S. H., and Prusiner, S. B. (2018b). Msa prions exhibit remarkable stability and resistance to inactivation. Acta Neuropathologica, 135(1):49–63.

Woerman, A. L., Oehler, A., Kazmi, S. A., Lee, J., Halliday, G. M., Middleton, L. T., Gentleman, S. M., Mordes, D. A., Spina, S., Grinberg, L. T., Olson, S. H., and Prusiner, S. B. (2019). Multiple system atrophy prions retain strain specificity after serial propagation in two different tg(snca*a53t) mouse lines. Acta Neuropathologica, 137(3):437–454.

Woerman, A. L., Patel, S., Kazmi, S. A., Oehler, A., Lee, J., Mordes, D. A., Olson, S. H., and Prusiner, S. B. (2020). Kinetics of α-synuclein prions preceding neuropathological inclusions in multiple system atrophy. PLOS Pathogens, 16(2):e1008222.

Woerman, A. L., Stöhr, J., Aoyagi, A., Rampersaud, R., Krejciova, Z., Watts, J. C., Ohyama, T., Patel, S., Widjaja, K., Oehler, A., Sanders, D. W., Diamond, M. I., Seeley, W. W., Middleton, L. T., Gentleman, S. M., Mordes, D. A., Südhof, T. C., Giles, K., and Prusiner, S. B. (2015). Propagation of prions causing synucleinopathies in cultured cells. Proc Natl Acad Sci U S A, 112(35):E4949–58.

Yamasaki, T. R., Holmes, B. B., Furman, J. L., Dhavale, D. D., Su, B. W., Song, E. S., Cairns, N. J., Kotzbauer, P. T., and Diamond, M. I. (2019). Parkinson’s disease and multiple system atrophy have distinct α-synuclein seed characteristics. J Biol Chem, 294(3):1045–1058.

Yang, Y., Shi, Y., Schweighauser, M., Zhang, X., Kotecha, A., Murzin, A. G., Garringer, H. J., Cullinane, P. W., Saito, Y., Foroud, T., et al. (2022). Structures of *α*-synuclein filaments from human brains with lewy pathology. Nature, 610(7933):791–795.

